# Ancient origin and high diversity of zymocin-like killer toxins in the budding yeast subphylum

**DOI:** 10.1101/2024.09.27.615389

**Authors:** Padraic G. Heneghan, Letal I. Salzberg, Eoin Ó Cinnéide, Jan A. Dewald, Christina E. Weinberg, Kenneth H. Wolfe

**Affiliations:** Conway Institute, School of Medicine, University College Dublin, Dublin 4, Ireland; Department of Life Sciences, Institute for Biochemistry, Leipzig University, Leipzig, Germany

## Abstract

Zymocin is a well-characterized killer toxin secreted by some strains of the yeast *Kluyveromyces lactis*. It acts by cleaving a specific tRNA in sensitive recipient cells. Zymocin is encoded by a killer plasmid or virus-like element (VLE), which is a linear DNA molecule located in the cytosol. We hypothesized that a tRNA-cleaving toxin similar to zymocin may have caused the three parallel changes to the nuclear genetic code that occurred during yeast evolution, in which the codon CUG became translated as Ser or Ala instead of Leu. However, zymocin-like toxins are rare – both among species, and among strains within a species – and only four toxins of this type have previously been discovered. Here, we identified 45 new zymocin-like toxin genes in Saccharomycotina, the budding yeast subphylum, using a novel bioinformatics strategy, and verified that many of them are toxic to *S. cerevisiae* when expressed. Some of the new toxin genes are located on cytosolic VLEs, whereas others are on VLE-derived DNA integrated into the nuclear genome. The toxins are extraordinarily diverse in sequence and show evidence of positive selection. Toxin genes were found in five taxonomic orders of budding yeasts, including two of the three orders that reassigned CUG codons, indicating that VLEs have been parasites of yeast species for at least 300 million years and that their existence pre-dates the genetic code changes.

## Introduction

In many budding yeast species (subphylum Saccharomycotina), some strains secrete killer toxins that inhibit the growth of other yeasts. There are many different types of killer toxin systems, including ones encoded by cytosolic DNAs, cytosolic RNAs, or chromosomal genes (1). One of the best-known yeast killer toxins is zymocin (also called γ-toxin), which is encoded by a cytosolic linear DNA molecule in *Kluyveromyces lactis* (2-4). Killer strains of *K. lactis* contain two cytosolic DNAs, pGKL1 and pGKL2. These molecules were initially called cytosolic linear plasmids (3, 5), but more recently they have been called virus-like elements (VLEs) to reflect their phylogenetic relatedness to viruses such as adenovirus and bacteriophages (6). pGKL1 and pGKL2 are small multi-copy linear double-stranded DNAs, capped with a terminal protein at each end. They are replicated and transcribed independently of the nuclear genome, by their own DNA and RNA polymerases. pGKL1 is a 9 kb ‘killer’ plasmid encoding the toxin zymocin. pGKL2 is a 13 kb ‘helper’ plasmid coding for the housekeeping functions (e.g. replication, transcription, and initiation of translation) necessary for the maintenance and expression of both plasmids. pGKL1 cannot survive without the functions provided by the helper plasmid pGKL2, whereas pGKL2 is autonomous.

Zymocin is a heterotrimeric molecule with three subunits (α, β, γ) of which the γ-subunit is the active moiety of the toxin. The γ-subunit is an anticodon nuclease that acts by cleaving the anticodon loop of tRNAs in sensitive cells, arresting their cell cycle. Zymocin’s primary target is tRNA-Glu(UUC) (7-9). As well as coding for the toxin γ-subunit, pGKL1 also has a gene coding for a chitin-binding protein (CBP) with a chitinase domain, which is a precursor from which the toxin α- and β-subunits are released by post-translational processing. Το target a victim cell, the γ- and α/β-subunits are linked by a cysteine-disulfide bridge and secreted from the killer cell. The secreted complex (called the zymocin holotoxin) binds to the chitin component of the cell wall of the victim cell, and the γ-subunit enters to cleave its target tRNA (3, 10-12) (Figure 1). To prevent self-targeting by the secreted toxin, pGKL1 also encodes a protein that confers immunity by binding to zymocin’s γ-subunit and inactivating it (13, 14).

**Figure 1.**
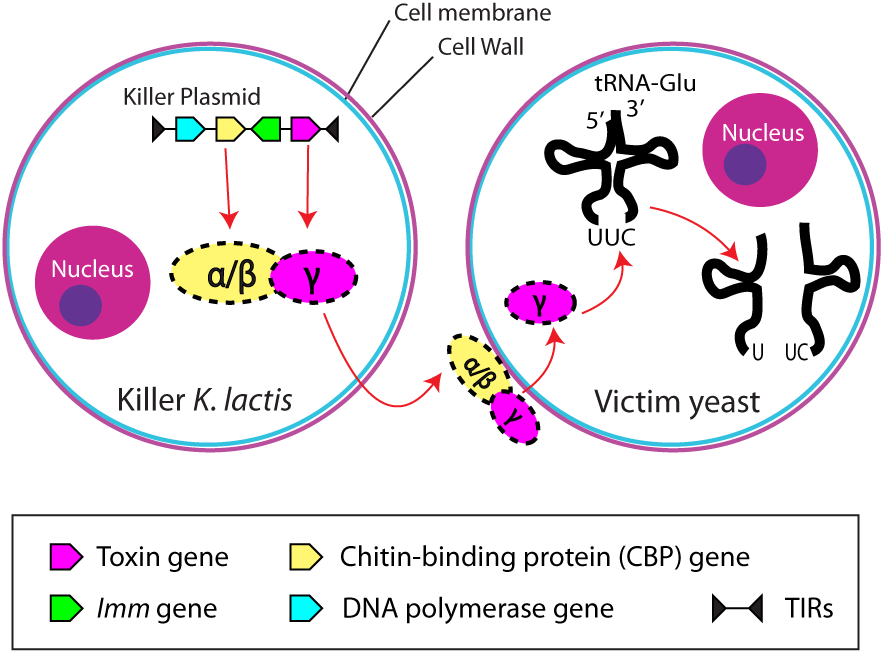
Summary of zymocin production and mechanism of action. A killer strain of *K. lactis* containing the pGKL1 killer plasmid produces and secretes a holotoxin consisting of chitin-binding (α/β) and ribonuclease toxin (γ) subunits. The secreted holotoxin attaches to the cell wall of a victim yeast cell, which can be a different species. The γ-subunit of the toxin translocates across the cell membrane and cleaves tRNA-Glu(UUC) molecules in the cytosol of the victim cell, stopping translation and killing it.

In the decades since zymocin was discovered, only three other VLE-encoded killer toxins have been found in budding yeasts. Two of them are anticodon nucleases with tRNA-Gln(UUG) as their major target: PaT from *Millerozyma acaciae* (formerly called *Pichia acaciae*; (15, 16)) and DrT from *Debaryomyces robertsiae* (17). The fourth toxin, PiT from *Babjeviella inositovora* (formerly called *Pichia inositovora*), acts by cleaving several sites in the 25S and 18S rRNAs (18) so technically it is not an anticodon nuclease because its target is not a tRNA. However, in all other respects PiT is functionally similar to the other toxins. Each of them is encoded by a killer plasmid, assisted by a helper plasmid, and the active γ-subunit is delivered to the recipient cell by a CBP.

VLEs are quite rare in yeasts. Many species, including *Saccharomyces cerevisiae*, do not contain them at all (19). Some species such as *Debaryomyces hansenii* and *Komagataella phaffii* (20, 21) have an autonomous helper plasmid and a smaller non-autonomous plasmid, but the smaller plasmid has no killing activity and does not appear to code for a toxin. These smaller plasmids are called ‘cryptic’ because their function is unknown. The most extensive search for VLEs was conducted by Fukuhara (5) who used gel electrophoresis to screen 1800 strains from 600 yeast species, discovering approximately 20 new VLEs. He found that only 1%-2% of strains have a VLE and that the presence of VLEs often differs among strains of the same species. However, none of the strains containing the new VLEs had clear killing activity against other yeasts, so the non-autonomous plasmids found by Fukuhara (5) appear to be cryptic plasmids, although they were not sequenced.

In this study we searched systematically for new zymocin-like toxin genes in budding yeast genome sequence data. Our motivation for this search was an hypothesis that we proposed in 2018 – that a zymocin-like anticodon nuclease killer toxin could have been the driver of the evolutionary reassignments of the CUG codon in the nuclear genetic code that occurred during budding yeast evolution (22). The codon CUG in nuclear genes is normally translated as leucine, but in three separate yeast lineages it was reassigned to either serine or alanine (22-26). These parallel genetic code changes occurred in the ancestors of three clades that were initially called the CUG-Ser1, CUG-Ser2 and CUG-Ala clades and are now recognized as the taxonomic orders Serinales, Ascoideales, and Alaninales (22, 27). Finding new candidate toxin genes is not straightforward because the level of sequence identity among the known toxins is very low. The anticodon nuclease (γ) subunits of PaT and DrT have 38% amino acid sequence identity to each other, but there is no statistically significant sequence similarity between this pair and either zymocin or PiT. Their lack of similarity raises the question of whether the known toxins are even orthologs (18, 28), but they are all encoded by VLEs and they all use the same mode of action: delivery of a ribonuclease to the recipient cell by a CBP. Here, we developed an innovative database search strategy and discovered a large and diverse set of potential new zymocin-like toxins, including both plasmid-encoded toxins and toxin genes located in nuclear genomes. We expressed many of the new toxin candidates in *S. cerevisiae* and found that they abolish or inhibit growth. Our results show that zymocin-like anticodon nuclease killer toxins have a much wider phylogenetic distribution, and an even higher level of sequence diversity, than previously recognized.

## Results

### Candidate toxin genes occur throughout Saccharomycotina, in VLEs and in nuclear genomes

We used multiple databases and search strategies to find new toxin candidates. The databases we searched included all the genome assemblies available in NCBI from budding yeast species (subphylum Saccharomycotina), including recent data for over 1000 species from the Y1000+ project (29, 30). We also made *de novo* genome assemblies (31) of 1375 Saccharomycotina isolates (381 unique species) for which Illumina sequencing reads were available in NCBI’s Sequence Read Archive (SRA). We made these *de novo* assemblies for two reasons: (i) some of the SRA datasets come from population genomics studies in which individual genomes were sequenced but not assembled, but some individuals in a population may contain VLEs that are absent from the reference genome assembly for that species; and (ii) we were concerned that some researchers might have omitted VLEs from the genome assemblies they submitted to databases, due to their small contig sizes and sequence coverage different from the nuclear genome. In addition, we sequenced most of the plasmids that were detected in the electrophoresis survey by Fukuhara (5), by whole-genome Illumina sequencing of the corresponding strain from the Westerdijk Institute (CBS) collection. We used TBLASTN searches (32) as described in Methods to find VLEs and candidate killer toxin genes in these databases.

The phylogenetic distribution of the VLEs and candidate toxins we discovered is summarized in Figure 2, and examples of VLE structures are shown in Figure 3. We found 10 new VLEs coding for candidate killer toxins, in species in the orders Saccharomycetales, Ascoideales, Serinales, and Dipodascales. More surprisingly, we found a further 35 candidate toxin genes that are located in nuclear genomes (Figure 2). In almost all of these cases the nuclear toxin gene is beside genes (or pseudogenes) for CBP and/or candidate immunity proteins (Figure 3). These nuclear loci appear to be the result of integrations of VLEs into chromosomes (33, 34), and in one case (*Metschnikowia hibiscae*) the nuclear integration includes a terminal inverted repeat (TIR), clearly indicating a recent history as a free VLE. Many of the nuclear loci are subtelomeric, for example in *Kazachstania barnettii* where two of its three candidate toxin genes occur in clusters with CBP and immunity genes at the ends of chromosome-sized contigs; these toxin proteins have only 82% amino acid identity. Phylogenetically, the newly-discovered candidate killer toxins come from species distributed across Saccharomycotina. Toxins were previously known in two of the 12 taxonomic orders (Saccharomycetales and Serinales), but we found new toxin candidates in three more orders: Ascoideales (VLE-encoded), Pichiales (nuclear-encoded), and Dipodascales (both VLE- and nuclear-encoded) (Figure 2).

**Figure 2.**
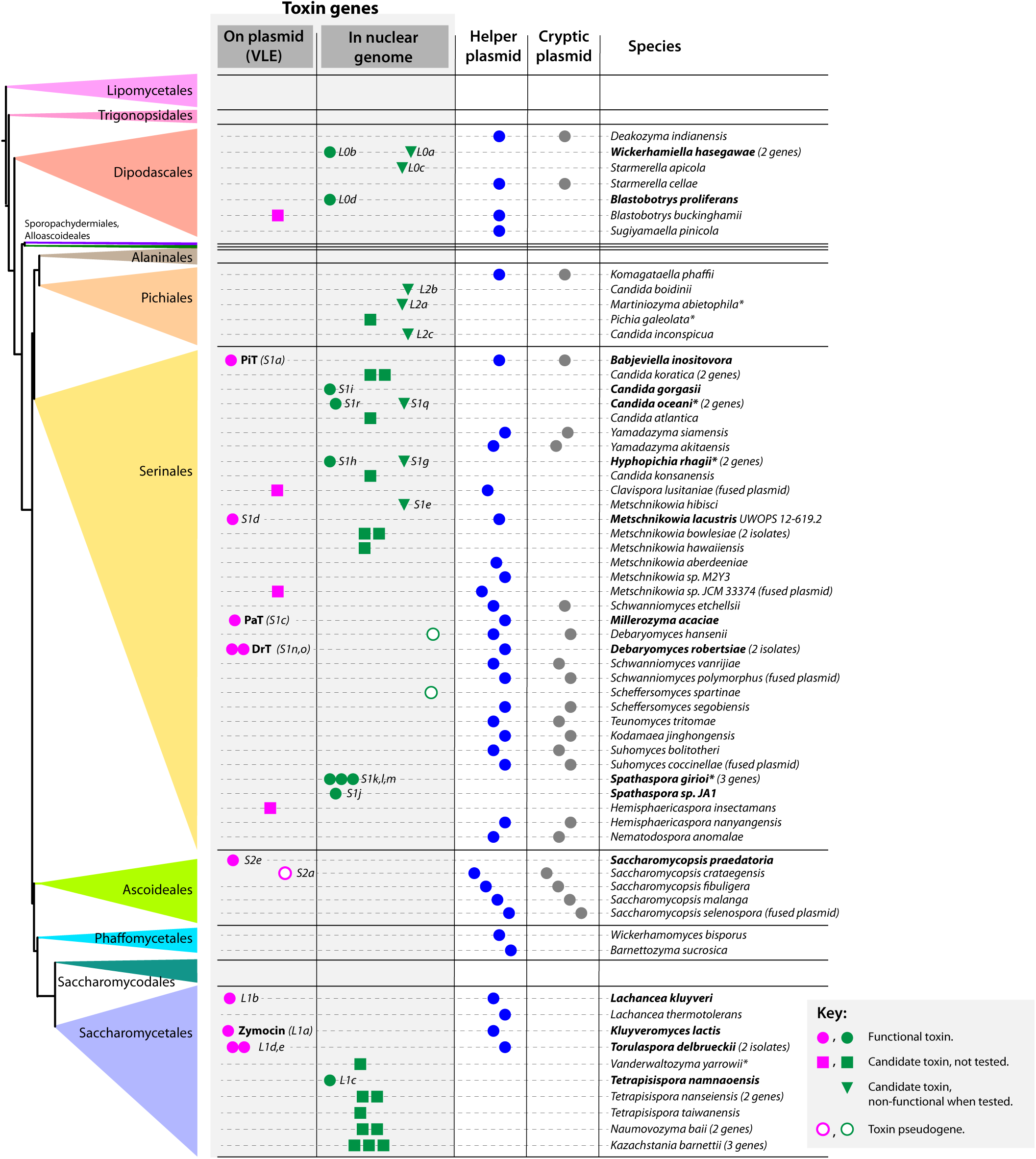
Phylogenetic distribution of toxin genes and cytosolic linear plasmids discovered in budding yeasts. The phylogenetic tree on the left shows the 12 taxonomic orders in the subphylum Saccharomycotina (27, 30). Species containing toxin genes that were functional in our assays are named in bold. Symbols in the four columns indicate candidate toxin genes located on linear plasmids (VLEs; pink), candidate toxin genes located in the nuclear genome (green), species containing helper plasmids (blue), and species containing cryptic plasmids (gray). For candidate toxin genes, symbol shapes and locations indicate ones that were functional when assayed in *S. cerevisiae* (filled circles), non-functional when assayed in *S. cerevisiae* (triangles), not tested (squares), or pseudogenes (open circles). Toxin candidates that were assayed were given names such as L0a, indicating the clade (L0, L1, L2, S1 or S2) followed by a lowercase letter. Some species containing only toxin pseudogenes or toxin genes with incomplete sequences have been omitted to save space. Asterisks beside species names indicate genome assemblies in which the toxin gene is on a short contig that lacks identifiable nuclear genes, but we scored them as nuclear contigs because we did not find helper plasmids in these species.

**Figure 3.**
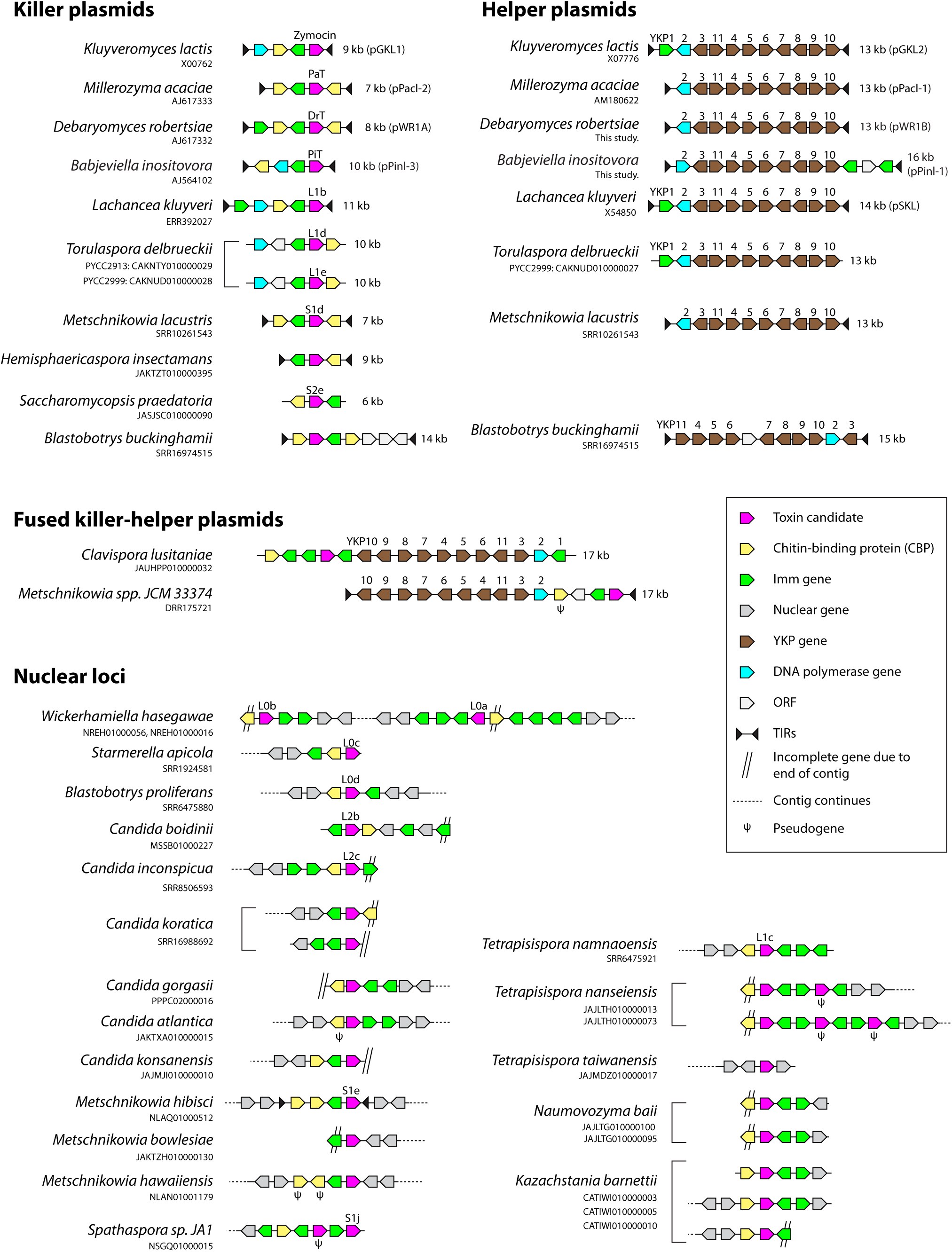
Examples of gene maps of VLEs and nuclear loci encoding candidate toxins. The killer plasmids coding for the four previously-known toxins zymocin, PaT, DrT and PiT are shown at the top for comparison to the newly-discovered loci. Names are shown for toxins that we assayed functionally. Genes in helper plasmids are named *YKP1* to *YKP11*, using the nomenclature introduced by Love et al. (62). Accession numbers from the NCBI nucleotide or SRA databases are shown. Not drawn to scale.

We found helper plasmids in 36 species, and cryptic plasmids in 21 species (Figure 2). In almost all cases where a yeast strain contains a killer plasmid (i.e., a VLE that codes for a toxin candidate) or a cryptic plasmid, this non-autonomous plasmid was found to be accompanied by a helper plasmid. Most of the helper plasmids are almost identical in terms of gene content and gene order, whereas the killer plasmids are much more variable, with several containing multiple CBP or immunity genes (Figure 3). We found five instances where fusions have occurred between a helper plasmid and either a killer plasmid or a cryptic plasmid, to form large autonomous linear plasmids up to 22 kb in size (the two killer-helper fusion plasmids are shown in Figure 3). We constructed a phylogenetic tree of helper plasmids from concatenated alignments of the conserved set of proteins they encode (Figure S1). Its topology is similar to the phylogeny of the host species, so there is little indication that VLEs have been exchanged extensively by horizontal transfer among yeast species.

### Many candidate toxin genes are functional in *S. cerevisiae*

To investigate whether the candidate genes code for functional toxins, we integrated them into *S. cerevisiae* under the control of an inducible promoter, after removing the secretion signal (replacing it with a start codon) and optimizing codon usage. Because the candidate protein is not secreted, and because no immunity proteins are present, functional toxins are expected to lead to cell death after induction if they can cleave an *S. cerevisiae* tRNA. For most toxins, we used a system in which the heterologous gene is induced using the hormone β-estradiol (35, 36). Examples of assays of three functional toxins are shown in Figure 4. In each case, the presence of β-estradiol dramatically reduces growth, on both solid and liquid media. We were unable to clone some toxin candidates into the β-estradiol system, probably due to leaky expression, so we assayed them instead by using induction by galactose from the *pGAL1* promoter. We verified that all four previously known toxins (zymocin γ-toxin, PaT, DrT and PiT) are functional in our assays, although we found that the sequences originally published for the DrT and PiT genes (37, 38) contain frameshift errors that render them non-functional. By re-sequencing the DrT and PiT genes, and expressing the correct sequences in *S. cerevisiae*, we confirmed that they encode functional toxins (Figure S2).

**Figure 4.**
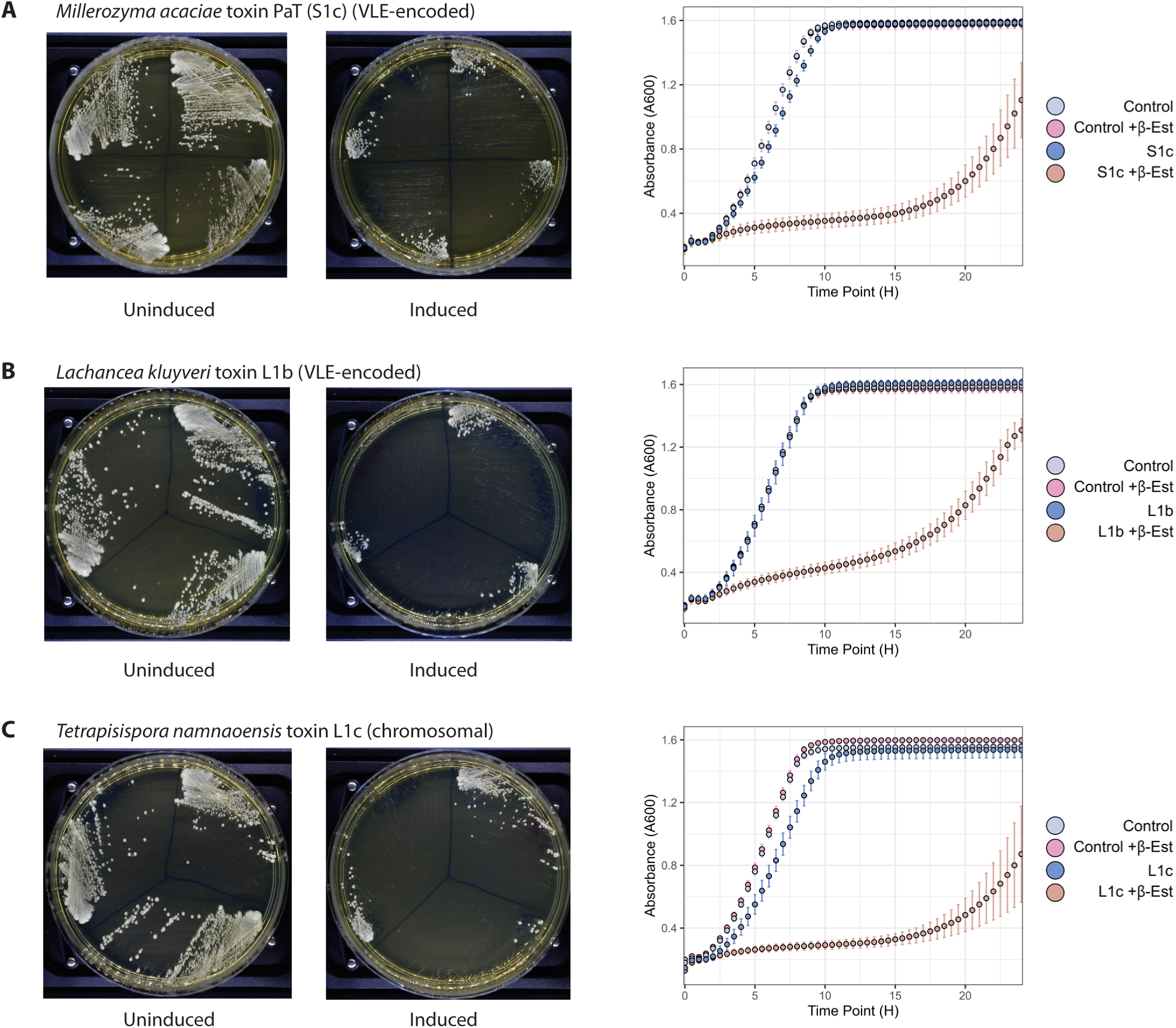
Examples of functional assays of toxin candidates: (A) the known toxin PaT encoded by a VLE in *Millerozyma acaciae*; (B) the newly discovered toxin L1b encoded by a VLE in *Lachancea kluyveri*; and (C) the newly discovered toxin L1c encoded by a nuclear gene in *Tetrapisispora namnaoensis*. Each toxin gene, without its secretion signal, was cloned into *S. cerevisiae* under the control of a β-estradiol inducible promoter. Plate images show 3 or 4 independent clones streaked onto YPD agar (left plates) or YPD + 2 μM β-estradiol (right plates), photographed after 48 hours. Graphs show growth curves of the same strains in liquid YPD media with or without 2 μM β-estradiol, for the tested toxin candidates and an *S. cerevisiae* control strain (PHY040) that contains the β-estradiol induction system but has no gene inserted downstream of the inducible promoter.

In total we assayed 24 new toxin candidates and found that 16 of them (67%) abolish or significantly reduce growth when induced in *S. cerevisiae*. Assays of all the tested toxin candidates are shown in Figure 4 and Figure S2, and summarized in Table S1. We found six new functional VLE-encoded toxins from three taxonomic orders: Saccharomycetales (toxins named L1b, L1d, L1e), Ascoideales (S2e), and Serinales (S1d, S1n) (Figure 2). We tested nuclear genes for candidate toxins and found 10 that are functional against *S. cerevisiae*, from three orders: Saccharomycetales (L1c), Serinales (S1h, S1i, S1j, S1k, S1l, S1m, S1r) and Dipodascales (L0b, L0d) (Figure 2; Table S1). In several species, a single nuclear locus contains two genes coding for candidate toxins with different sequences, and in each pair that we tested we found that one of the genes was toxic when induced in *S. cerevisiae*, and the other was not (L0b/L0a, S1r/S1q, and S1h/S1g; Figure 2).

### Identification of targets of *Lachancea kluyveri* toxin L1b by cyPhyRNA-seq

We characterized a new toxin, L1b from *Lachancea kluyveri*, in more detail. A cytosolic linear helper plasmid from *L. kluyveri* was identified many years ago and named pSKL (39). It was discovered in strain IFO 1685, but its function remained unclear because IFO 1685 does not contain a killer plasmid and has no killer activity. By assembling Illumina data from SRA for 28 isolates of *L. kluyveri* that were sequenced for a population genomics study (40), we detected the helper plasmid pSKL in three isolates. One of these isolates, NCYC 543, also contains a killer plasmid. NCYC 543 should be identical to IFO 1685 because they are both culture collection deposits of the type strain of *L. kluyveri*, but NCYC 543 has retained a killer plasmid that IFO 1685 has lost. Both of these plasmids are absent in SRA data from CBS 3082, a third culture collection deposit of the type strain, suggesting that they are lost easily. The killer plasmid encodes a functional toxin that we named L1b (Figure 3; Figure 4B). The closest relatives of L1b are two newly discovered toxins from *Torulaspora delbrueckii* (35% and 43% amino acid identity), and *K. lactis* zymocin (28%).

We used a recently developed method, cyPhyRNA-seq (41), to characterize the tRNA targets of L1b and, as a control, the canonical toxin PaT. When anticodon nuclease ribotoxins cleave their target tRNAs, they leave an unusual 2’,3’ cyclic phosphate group (instead of 3’-OH) at the end of the upstream fragment (7, 16). In cyPhyRNA-seq, libraries for high-throughput sequencing are constructed by specifically ligating adapters onto small RNAs that have 2’,3’ cyclic phosphate ends. We sequenced cyPhyRNA-seq libraries from *S. cerevisiae* cultures in which expression of either PaT or L1b was induced by addition of β-estradiol, and compared them to uninduced controls. For PaT, we detected specific enrichment of tRNA fragments terminating in the anticodon loops of tRNA-Gln(UUG) and tRNA-Gln(CUG), which are known to be its major and minor targets respectively (Figure S3A) (15, 16). For L1b, we detected specific enrichment of tRNA fragments terminating in the anticodon loop of tRNA-Glu(UUC), which is the major target of zymocin, and in the variable loop of tRNA-Ser(IGA) (Figure S3B). Cleavage of the variable loop of a tRNA by a toxin has not been reported before, and further experiments will be required to validate these putative targets of L1b. We transformed plasmids expressing tRNA-Glu(UUC) and/or tRNA-Ser(IGA) into the host strains, but they did not rescue the growth impairment caused by induction of L1b.

### Rapid evolution and positive selection on toxin genes

The toxin sequences are evolving rapidly and are diverse even within a single species. For example, we identified two similar VLEs from two isolates of *Torulaspora delbrueckii* that code for toxins with only 35% amino acid identity to each other (L1d and L1e; Figure 3), whereas two isolates of *Debaryomyces robertsiae* contain VLEs coding for variants of DrT with 90% identity. For species with nuclear genes coding for two candidate toxins, the sequence identity of the two toxins ranges from no significant similarity (*Hyphopichia rhagii*) to 35% (*Wickerhamiella hasegawae*) to 91% (*Tetrapisispora nanseiensis*).

It is not possible to construct a reliable phylogenetic tree of the toxins due to the low sequence identity among many of them. Instead, we used BLASTP (32) and Cytoscape (42) to represent their sequence relationships as networks (Figure 5). In this diagram, two toxin proteins (nodes) are connected by an edge if the BLASTP hit between them exceeds marginal significance (E-value < 1e-6). Four clusters of related toxins (designated Tox-A to Tox-D) are apparent, as well as two other toxin pairs and six singletons (Figure 5), making a total of 12 groups of toxin sequences that are essentially unrelated to one another. The Tox-A cluster consists of all the toxins from Saccharomycetales, including zymocin, and all but one of the toxins from Pichiales. The greatest diversity occurs in Serinales species, whose toxins lie in five sequence groups (Tox-B, Tox-C, Tox-D and two singletons). The toxin candidates identified in other orders (Dipodascales and Ascoideales) are all singletons or only have relatives within their own genus. At least eight different sequence groups contain functional toxins (Tox-A, Tox-B, Tox-C, and five others; Figure 5). Interestingly, neither of the toxin candidates that we tested from the Tox-D cluster had activity against *S. cerevisiae* in our assays.

**Figure 5.**
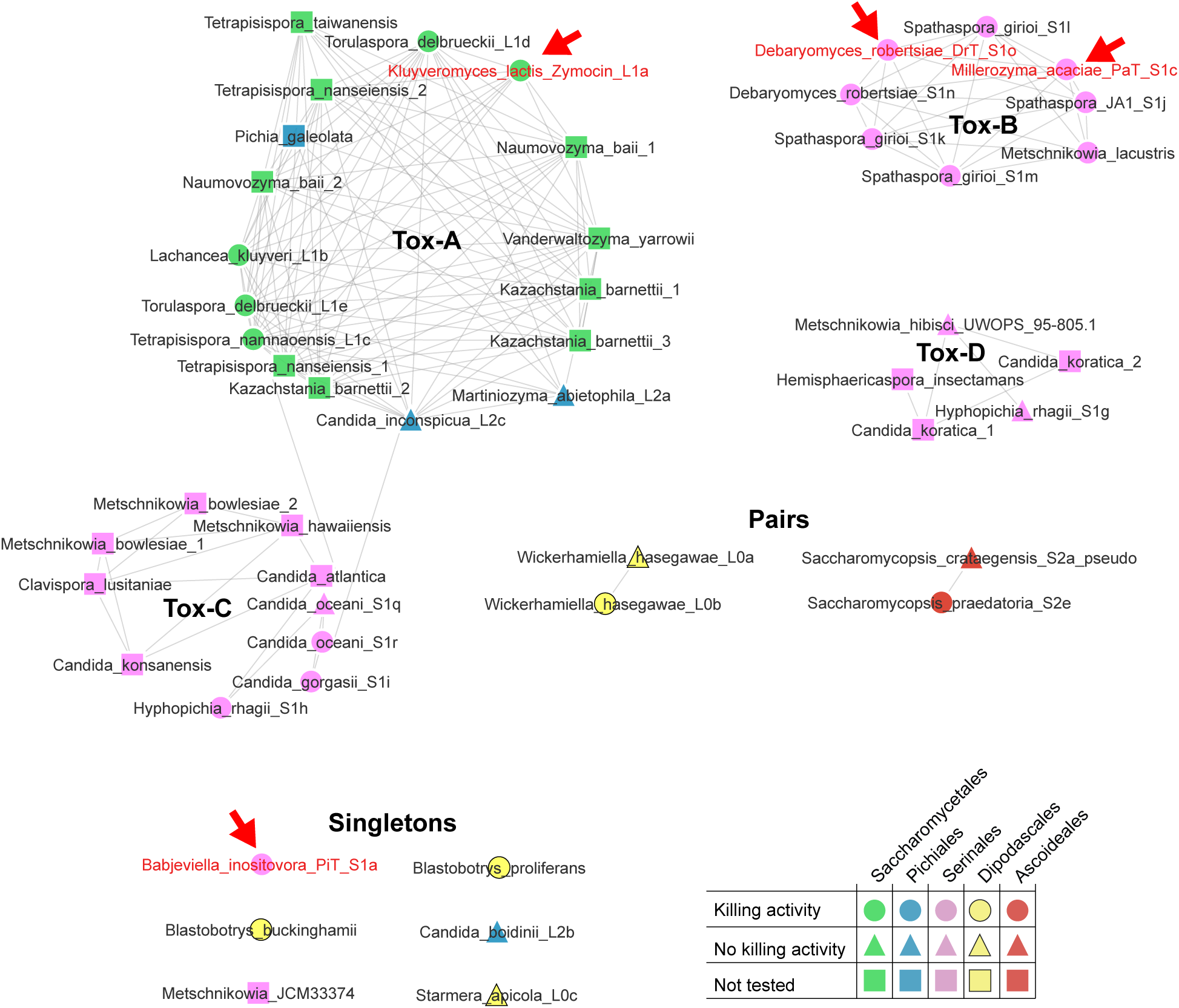
Rapid evolution of toxins. Each node represents a toxin protein. Nodes are connected by edges if they share even borderline amino acid sequence relatedness (BLASTP *E*-value < 1e-6), with thicker edges indicating greater significance (lower *E*-values). Node colors and shapes indicate their taxonomic order and functional activity in our assays as shown in the key. Red arrows indicate the four canonical toxins.

The high sequence diversity of the toxin proteins suggested that they might be evolving under positive (Darwinian) selection. Analysis of the nonsynonymous-to-synonymous nucleotide substitution ratio in pairs of toxin genes from the same species showed that overall, the genes are subject to purifying selection on their amino acid sequence (dN/dS ratios between 0.15 and 0.60 over the whole gene; ref. (43)). To test for positive selection on individual codon sites, we analyzed a set of seven toxin sequences from the Tox-B cluster that are all functional and relatively conserved in sequence. This group comprises PaT, DrT (two isolates), and four *Spathaspora* toxins. Amino acid sequence identity among these seven sequences ranges from 27% to 90%. Using CODEML (44), Naive Empirical Bayes and Bayes Empirical Bayes (BEB) analyses agreed on four residues that appear to be under positive selection (*P* > 0.95), while BEB identified two additional residues with *P* > 0.95 (Table S2). Four of these six residues surround the active site of the toxin ribonuclease (Figure S4). Interestingly, one of the two positively selected residues identified by BEB, Asn-116, lies within one of the toxin’s 3_10_ helices that surround the active site and have high positive potential, implying a role in RNA binding (16). Finally, one of the residues shown to be essential for toxin activity in DrT and PaT (16), Lys-175, is not conserved in the *Spathaspora* toxins which indicates that essential sites can be specific only to close groups of toxins.

### Diversity and multiplicity of immunity-related genes

The VLEs that code for zymocin, PaT and DrT also contain genes coding for non-secreted proteins that have antitoxin activity, giving the killer cell immunity to the toxin it secretes. We refer to these three genes as the canonical immunity genes because their role in immunity has been demonstrated experimentally (13, 14, 17, 45). Each of the three canonical immunity genes is located immediately adjacent to the gene for the toxin that it counteracts (Figure 3). The canonical immunity genes also have a large and diverse family of homologs, which we call *Imm* genes because we postulate that they have an immunity function too, although this has not been demonstrated experimentally. *Imm* genes are present on many toxin-encoding VLEs, sometimes with multiple paralogous copies (Figure 3). *Imm* genes are also present on some helper plasmids (e.g. *ORF1/YKP1* of *K. lactis* pGKL2) and cryptic plasmids, and beside putative nuclear toxin genes (Figure 3). In addition, there are *Imm* genes in the nuclear genomes of many yeast species in which we did not identify a nuclear toxin gene or a VLE.

Imm proteins are diverse, but less diverse than the toxins. We used BLASTP and Cytoscape to explore the relationships among Imm proteins from the species in which we had identified candidate toxins, using a relatively stringent BLASTP threshold of E < 1e-22 to connect nodes and build clusters (Figure 6); at more liberal thresholds, these clusters merge and all the Imm proteins are related to each other. As with the toxins (Figure 5), the Imm proteins show a degree of clustering into phylogenetic groups, and high sequence diversity within the Serinales.

**Figure 6.**
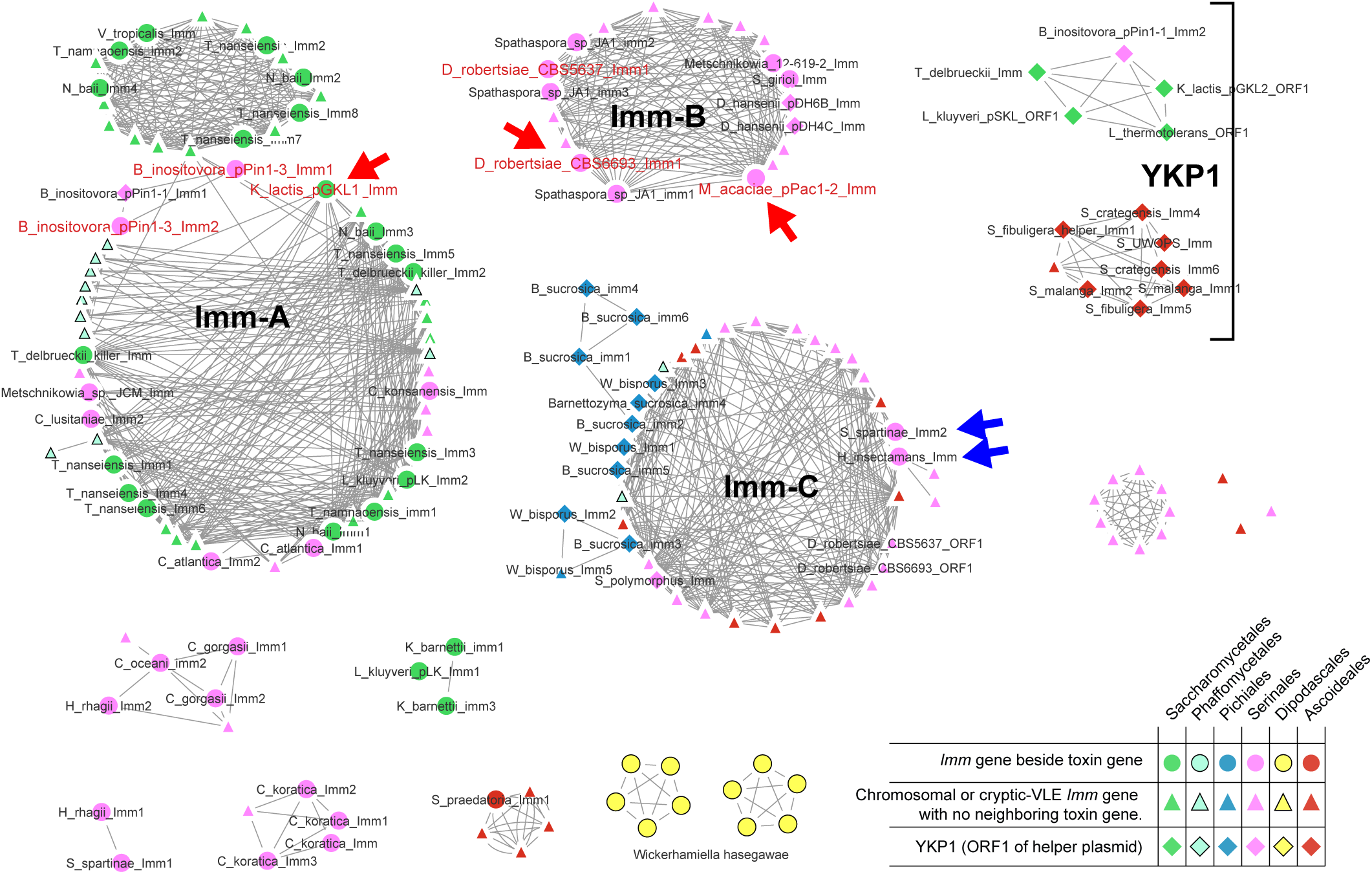
Sequence clusters of predicted immunity (Imm) proteins. Nodes are connected by edges if they share strong amino acid sequence relatedness (BLASTP *E*-value < 1e-22), with thicker edges indicating greater significance (lower *E*-values). Node colors indicate their taxonomic order, and node shapes indicate the location of the *Imm* gene, as shown in the key. *Imm* genes located beside toxin genes (circles), and *Imm* genes located on helper plasmids (diamonds) are identified by name. Red arrows indicate the experimentally characterized immunity genes from *K. lactis* (pGKL1 ORF3), *M. acaciae* (pPacl-2 ORF4), and *D. robertsiae* (pWR1A ORF5). Three large clusters are labeled Imm-A, Imm-B and Imm-C. The label YKP1 marks two clusters of *Imm* genes that are located on helper plasmids, including *K. lactis* pGKL2 ORF1. Blue arrows highlight the only two *Imm* genes in the Imm-C cluster that are located beside toxin genes.

There are three distinct large clusters of Imm proteins. Two of them (Imm-A and Imm-B) include the canonical immunity proteins associated with the canonical killer toxins (red arrows in Figure 6), as well as uncharacterized *Imm* genes that are located on VLEs immediately beside candidate toxin genes (indicated by circular nodes). The third large cluster (Imm-C) consists of *Imm* genes that are mostly not located beside toxin genes; many of them are chromosomal or on cryptic plasmids, and there are only two toxin-adjacent *Imm* genes in this large cluster. The Imm-C cluster also includes ORF1 of the *D. robertsiae* killer plasmid pWR1A, an *Imm* gene whose function is unknown; it is distinct from the canonical pWR1A immunity gene *ImmDrT (*pWR1A ORF5, which is in the Imm-B cluster) and not located beside the DrT toxin gene. Since the *Imm* genes in the Imm-C cluster are mostly not neighbors of toxin genes, we suggest that they may be genes conferring immunity to toxins that their hosts do not encode, but sometimes encounter.

Positive selection analyses of the canonical immunity genes in the Imm-B cluster and their close relatives indicated that three residues are under positive selection (*P* > 0.99; agreement between BEB and NEB) (Table S3; Figure S5). Considering that the characterized immunity proteins ImmDrT and ImmPaT have higher sequence identity (50%) than their toxins DrT and PaT (38%), and that ImmPaT confers some cross-immunity to DrT whereas ImmDrT does not confer cross-immunity to PaT (17), the immunity proteins may have only a few residues that confer complete immunity to a specific toxin. The positively selected residues may assist with stabilizing the interactions between the immunity protein and its cognate toxin.

## Discussion

Our work has greatly increased the number of zymocin-like toxins known, beyond the canonical four. More than half of the new toxin genes we tested were functional in our assays; we found six new functional toxins encoded on VLEs, and 10 encoded by chromosomal genes (Figure 2). The candidate toxin genes that were nonfunctional in our assays may be pseudogenes, but it is also possible that they encode toxins whose natural targets are tRNAs (or other RNAs) in yeast species other than *S. cerevisiae* – in other words, that they are toxins whose targets are absent in *S. cerevisiae* or are too divergent to be recognized and cleaved. We found that VLEs frequently integrate into yeast nuclear genomes, and in many cases the toxin proteins encoded by these integrations are functional. Whether these integrations represent successful functional relocations of VLE genes to the nucleus, overcoming the barrier of A+T richness and promoter incompatibility (14), or simply the result of recent nuclear capture of VLE DNA that has not yet degenerated (NUPAVs (33, 34)) is unclear.

We detected candidate toxin genes (either VLE-encoded or chromosomal) in 14 (1.0%) of the 1375 genome assemblies we made from the SRA database, and in 45 (1.2%) of the 3780 assemblies used from NCBI. These frequencies are consistent with Fukuhara’s observation that 1-2% of yeast isolates contain VLEs of any type (5). In total, we found toxin candidates in five of the 12 taxonomic orders of Saccharomycetales, and helper plasmids in a sixth (Figure 2). The orders in which we did not find toxins are relatively small, represented by fewer species and genome sequences, so in view of the overall rarity of toxins the failure to find them in these orders is not surprising. Since the phylogenetic tree of helper plasmids indicates that VLEs are largely inherited vertically (Figure S1), the results indicate that although VLEs are rare, they are old. Zymocin-like anticodon nucleases can therefore be inferred to have been present through most or all of the evolutionary history of the whole subphylum Saccharomycotina. Their minimum age corresponds to the date of the last common ancestor of the 10 orders excluding Lipomycetales and Trigonopsidales (Figure 2), which has been estimated to be 326 Myr (29). Anticodon nucleases therefore pre-date the emergence of the orders with altered genetic codes (Serinales, Alaninales and Ascoideales), which is consistent with the hypothesis that a killer toxin may have been a driver of the CUG codon reassignment (22). However, no toxin targeting tRNA-Leu(CAG) has yet been identified, and since this tRNA isotype is not present in *S. cerevisiae*, it cannot be the sole target of any of the new toxins we found to be functional in *S. cerevisiae*. Identifying the tRNA targets of the new toxins, and investigating how toxins coevolve with their tRNA targets and with immunity proteins, will be a challenge for future studies.

We found that VLE structures are fluid. Although most helper plasmids are almost identical in gene organization (Figure 3), we found five separate instances where a helper plasmid has become fused to either a killer plasmid or a cryptic plasmid, forming VLEs up to 22 kb in size. We also found many cryptic plasmids throughout the Saccharomycotina. The functions of these plasmids remain unclear; they are non-autonomous plasmids lacking an identifiable γ-toxin gene, but most of them contain genes for one or more secreted chitin-binding proteins related to toxin α/β subunits.

Why are toxin proteins so diverse? We suggest that they may be engaged in two different evolutionary competitions. First, they may be in competition with tRNAs. Mutations in a tRNA gene that enable it to evade a toxin will be favored, but so too will mutations in a toxin that restore its ability to recognize either its original tRNA target or a different tRNA. Second, toxins may be in competition with immunity proteins, which can be encoded by the same VLE, by other VLEs, or by the nuclear genome of the target yeast species. Our finding that some sites in both toxin genes and immunity genes are under positive selection is consistent with coevolution of these two groups of genes.

It is interesting that multiple paralogous *Imm* genes are sometimes present in the same VLE, and very often present at the same nuclear locus (Figure 3). The highest numbers that we found were 6 *Imm* genes in a helper plasmid VLE of *Barnettozyma sucrosica*, and 16 in a nuclear locus of *Kazachstania martiniae*. The most plausible reason why a yeast cell would maintain multiple different immunity genes, while encoding only one toxin or even no toxin, is that it regularly encounters a variety of toxins, secreted by other cells, that it needs to overcome. We suggest that yeast cells may be under selection to maintain a defensive array of immunity genes to protect them from exogenous zymocin–like toxins. This exogenous immunity is separate from the need for a yeast cell carrying a toxin gene (on a killer plasmid or in its nuclear genome) to also carry an immunity gene that specifically protects it from this endogenous toxin.

Selection for immunity to diverse exogenous zymocin-like toxins provides a possible explanation for the frequent finding of yeast isolates that contain helper plasmids but not killer plasmids, and for the persistence of cryptic plasmids in populations, because helper plasmids and cryptic plasmids often contain *Imm* genes. Indeed, it is even possible that the CBPs encoded by cryptic plasmids serve a defensive function, cloaking the cell wall of the cell containing the plasmid, rather than an offensive function. A similar mechanism is used by pathogenic Pezizomycotina fungi, which coat themselves in LysM proteins during infections of insects and plants to protect themselves from chitinases secreted by the host (46, 47). It is possible that cryptic plasmids act in an analogous fashion, in which case they should be regarded as defense elements.

## Materials and Methods

### Bioinformatics Methods

We carried out searches for VLEs and toxin genes in the budding yeast (subphylum Saccharomycotina) genome sequence data available from the U.S. National Center for Biotechnology Information (NCBI) during 2023, either as assemblies deposited in the nucleotide database (3780 assemblies from 1122 species), or as unassembled reads deposited in the SRA (Sequence Read Archive) database (1375 SRA datasets from 381 species). The SRA datasets were downloaded using SRA Explorer (https://sra-explorer.info/), trimmed using Skewer v0.2.2 (48), and assembled using SPAdes v3.14 (31). We also searched our assemblies of the genomes that were newly sequenced for this study.

As well as using the four previously known killer toxin proteins as queries in TBLASTN searches, we also used CBP (chitin-binding protein), Imm (immunity-like), and DNA polymerase proteins from VLEs as queries because they evolve more slowly than toxins and are therefore more likely to retain significant sequence similarity. We also used YKP proteins from helper plasmids, including YKP6 (RNA polymerase) as TBLASTN queries. Contigs that were hit by any of these TBLASTN queries were annotated manually. Candidate killer toxin genes were identified as open reading frames adjacent to *CBP* or *Imm* genes, coding for proteins with a predicted secretion signal (TargetP score > 0.9; (49)), three conserved amino acid residues corresponding to Glu-9, His-287 and Cys-317 of PaT (28), and a total length of 200-300 amino acids. When new toxin candidates were found, they in turn were used as queries for further TBLASTN searches.

A phylogenetic tree of helper plasmids (Figure S1) was made by aligning individual homologous proteins (YKP2 to YKP11) using Muscle (50), concatenating the alignments, trimming using trimAl’s heuristic model (51), and then using IQ-Tree and ModelFinder for maximum-likelihood tree building (52, 53). Bootstrapping (1000 replicates) was executed using both UFBoot and SH-aLRT (54, 55). Visualization of sequence relationships among toxin proteins (Figure 5) and among immunity-like proteins (Figure 6) was done using Cytoscape (42). Tests for positive selection on toxin and *Imm* genes were performed using site model analysis in CODEML (44).

### New Genome Sequencing

We obtained 14 yeast strains from the Westerdijk Institute (CBS) culture collection that were reported by Fukuhara (5) to contain VLEs that have not subsequently been characterized, and cultured them at 25°C or 30°C as appropriate (Table S4). Using standard protocols (56), genomic DNA was extracted, sequenced on an Illumina NextSeq 550 instrument, and assembled using SPAdes v3.14 (31). We found that most of the VLEs in these assemblies were helper or cryptic plasmids (Table S4), but this experiment yielded a corrected sequence (designated S1o) for toxin DrT encoded by killer plasmid pWR1A from *Debaryomyces robertsiae* isolate CBS6693 (correcting the sequence originally reported for DrT (38)), and a second *D. robertsiae* isolate CBS5637 containing a pWR1A-like plasmid coding for a DrT-like toxin designated S1n that has 10% amino acid sequence divergence from S1o. We found that one of the three VLEs reported in *Saccharomycopsis crataegensis*, pScrl-3 (5, 57), is a killer-like plasmid but its candidate toxin gene (named S2a) was a pseudogene in all the *S. crataegensis* isolates we examined. We also sequenced *Babjeviella inositovora* isolate NRRL Y-12698 to obtain complete sequences of its three plasmids pPinl-1 (helper), pPinl-2 (cryptic) and pPinl-3 (killer). Our sequence of pPinl-3 includes corrections to the amino acid sequence of the *B. inositovora* toxin PiT (S1a) from this strain relative to its original publication (37).

### Characterization of Toxin Candidates

All toxin-expressing strains were constructed in *S*. *cerevisiae* strain PHY039 (Table S5), a derivative of strain MOY007 in which *HO* was deleted using a two-plasmid CRISPR-Cas9 system plus linear repair templates. MOY007 (Table S5) was constructed by transforming strain BY4742 (*MAT*α *his*3Δ1 *leu2*Δ0 *lys2*Δ0 *ura3*Δ0), with *Pac*I-linearized plasmid FRP880 (Table S6) and the Cas9-NAT plasmid (https://www.addgene.org/64329/) using a lithium acetate transformation protocol (58). Transformants were selected on histidine dropout medium plus 100 μg/mL nourseothricin (NTC; HKI Jena). This strain contains the fusion transcription factor LexA-ER-B112 (35) at the *HIS3* locus. PHY039 also contains an intronless *tL(CAG)* from *L. kluyveri* in place of the *S. cerevisiae tL(UAG)L2* gene, but this genotype was not utilized in the current study.

Among the candidate toxin genes that we identified, we chose 29 for expression in *S. cerevisiae*, including the canonical toxins zymocin, PaT, DrT and PiT (Table S1). Candidate toxin genes, without their predicted secretion signal sequences, were synthesized with codon optimization for *S. cerevisiae* by Twist Bioscience, and cloned into the integrating plasmid pRG634, which contains a β-estradiol inducible expression system (gift from Robert Gnügge) (36). Toxin-pRG634 vectors were linearized with *Asc*I and transformed into *S. cerevisiae* strain PHY039 using a lithium acetate transformation protocol (58) with selection on leucine dropout agar. PCR with internal primers was used to check for the presence of the toxin gene, and pRG634 junctions were amplified to check for correct insertion into the *LEU2* locus. At least three biological replicates of each toxin-expressing strain (except *Starmerella apicola* candidate L0c) were stocked in glycerol.

To test β-estradiol induction of candidate toxin genes on solid media, *S. cerevisiae* strains containing each toxin gene were streaked on YPD agar ± 2 μM β-estradiol as previously described (36), and photographed after 48 h incubation at 30° C. For liquid assays, overnight cultures of these strains were diluted to an OD600 of 0.1 in 200 μL of YPD or YPD + β-estradiol (0.5 μM) in 96-well plates, with duplicates for each biological replicate in each media (no difference was seen between 2 μM and 0.5 μM β-estradiol inductions of a positive control). OD600 was measured every 30 minutes for a 24 h period using a Synergy H1 microplate reader (BioTek).

To express toxins under galactose induction, the *GAL1* promoter was cloned into *Not*I/*Spe*I-digested toxin-pRG634 vectors (this removes the *LexO-CYC1* promoter). The resulting plasmids were linearized with *Asc*I and transformed into *S. cerevisiae* strain PHY039 with selection on leucine dropout agar. For galactose induction of these genes, overnight cultures grown in raffinose were washed twice in PBS and then each strain was diluted to an OD600 of 0.05 in 200 μL of SC+galactose (2%) media in a 96-well plate. OD600 was measured every 30 minutes for a 24 h period using a Synergy H1 microplate reader (BioTek).

### cyPhyRNA-sequencing

Cleavage of tRNAs by zymocin and PaT is known to leave 2’,3’ cyclic phosphate (2’,3’-cP) ends on the 3’-end of the 5’-tRNA fragment. Based on the similarity between the *Lachancea kluyveri* toxin L1b and zymocin, we hypothesized that L1b would make similar cleavage products. Thus, we performed cyPhyRNA-sequencing (41) to identify the targets of L1b. Briefly, PaT (positive control) and L1b-inducible strains were grown overnight in YPD, then back diluted 1:62.5 in 50 mL YPD (in duplicate; uninduced control/induced sample) the following day and grown at 30°C with shaking. After two doublings, one of the cultures was induced with 1 μM β-estradiol and the cultures were grown for another 2.5 h before harvesting the samples by centrifugation. The cell pellets were resuspended in 2 mL lyticase buffer (1 M sorbitol, 0.1 M EDTA, 0.1% β-mercaptoethanol) plus 50 U of lyticase (Sigma) and incubated for 30 min at 30°C. Total RNA was then extracted using the Monarch Total RNA Miniprep Kit (NEB). Subsequently, small RNA enrichment was performed using the mirVana miRNA Isolation Kit (Invitrogen) with 100 µg of total RNA. Libraries specific for RNAs with 2’,3’-cP (5’-tRNA fragments) were produced as described (41), except that the tRNA-blocking step was omitted. Libraries were purified using solid phase reverse immobilization beads (Vazyme VAHTS DNA Clean Beads) and sequenced with ∼30 million reads, 150 bp paired-end by Illumina HiSeq sequencing (Azenta Genewiz). Growth, lysis, and RNA extraction/enrichment were performed on the same day to prevent rapid degradation of the 2’,3’-cP moieties.

Reads from the cyPhyRNAseq Illumina libraries were processed using UMI-tools (59) to retain only unique reads and remove artifacts resulting from PCR amplification. After pre-processing with Skewer (48), reads were aligned using STAR (60) to a subset of the *S. cerevisiae* genome consisting of a single representative of each tRNA isodecoder, with gaps (N_10_) between each tRNA gene. Ratios of tRNA fragment terminus abundance between induced and uninduced cultures were analyzed using scripts adapted from Barth and Woychik (61). After alignment, 99% of positions (i.e., individual nucleotide sites in all tRNAs) in all libraries had abundances of ≤ 1 read per million, as expected given that tRNAs will generally fragment at specific positions, so these sites were removed by filtering. We then used the most abundant 15% of positions in the remaining data for heatmap analysis (Figure S3).

## Acknowledgments

This work was supported by the European Research Council (789341). We thank Robert Gnügge for strains and plasmids, Katherine Moreau for help with DNA sequencing, and Roy Walker for comments on the manuscript.

## Figure legends

**Figure S1.** Maximum-likelihood phylogenetic tree constructed from helper plasmid proteins YKP2-YKP11. The tree was rooted using *Deakozyma*. Colors indicate the taxonomic order of each species. Although the tree is not completely congruent with the species phylogeny (30) and there are some indications of paralogy or interspecies transfer (e.g. in the *Debaryomyces* species), the predominant pattern is one of vertical transmission.

**Figure S2.** Functional assays of toxin candidates on solid and liquid media. Each panel shows two plates, YPD uninduced (left) and YPD induced (right) for multiple independent clones of a toxin, and OD600 for growth in liquid media over 24 h with and without toxin induction. Strains were induced with either β-estradiol or galactose, as indicated. See Materials and Methods for details.

**Figure S3.** cyPhyRNA-seq investigation of tRNA targets of (A) the canonical toxin PaT from *Millerozyma acaciae*, and (B) the newly discovered toxin L1b from *Lachancea kluyveri*. Sequencing libraries were constructed using *Arabidopsis thaliana* tRNA ligase variant K152A/D726A to capture tRNA fragments having 2’,3’-cP termini, from *S. cerevisiae* strains in which expression of the toxin was either induced or uninduced. The Illumina sequencing reads were aligned to a database consisting of the DNA sequences of every tRNA isodecoder gene in *S. cerevisiae*, to map the tRNA fragment termini. The heatmaps (upper panels) show the locations of fragment termini that are enriched in the induced samples relative to the uninduced samples. Some termini are highlighted with arrows and names indicating the tRNA type and the nucleotide position of the detected terminus within it. The scatterplots (lower panels) illustrate the data processing steps that were applied to remove noise resulting from ratio calculations from low-abundance reads. They compare the ratio of reads in the induced to uninduced samples (Y-axis), to the abundance of the reads in the induced samples (X-axis; RPM, reads per million), for all tRNA fragment termini detected. We applied filters to remove tRNA fragment termini with <1 RPM, and show the ratios in the most abundant 15% of the remaining termini (points to the right of the dashed lines) in the heatmaps.

**Figure S4.** Residues of interest highlighted on the structure of PaT (ref. (16); PDB accession 4O87). Green, residues shown to be essential for toxin activity (16). Red, positively selected residues identified by both Naive Empirical Bayes (NEB) and Bayes Empirical Bayes (BEB) with *P* > 0.99. Orange, positively selected residues identified by both NEB and BEB with *P* > 0.95. Yellow, residues identified only by BEB as positively selected with *P* > 0.95. The positively selected sites N116, K119, S126, and V136 are located on the periphery of the active site, suggesting a role in stabilizing an interaction between a target and the active site.

**Figure S5.** Multiple sequence alignment of Imm proteins from the Imm-B cluster, with the positively selected sites identified by CODEML (Table S3) indicated. The three sites identified by both NEB and BEB with *P* > 0.99 (red dots) correspond to Q68, N220 and I239 of ImmPaT.

