## Supplementary Figures for "Ancient origin and high diversity of zymocin-like killer toxins in the budding yeast subphylum"

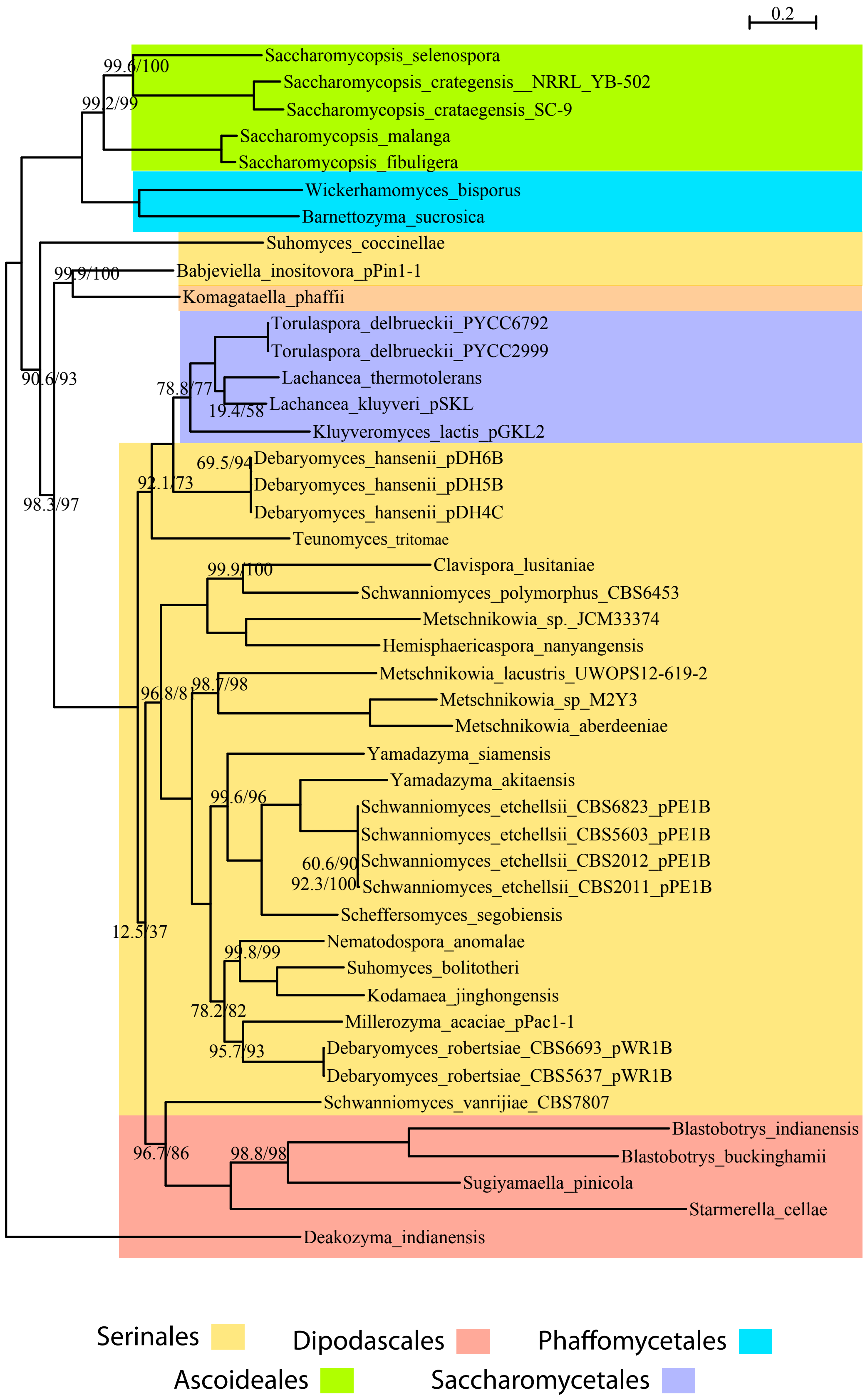


**Figure S1.** Maximum-likelihood phylogenetic tree constructed from helper plasmid proteins YKP2-YKP11. The tree was rooted using *Deakozyma*. Colors indicate the taxonomic order of each species. Although the tree is not completely congruent with the species phylogeny (30) and there are some indications of paralogy or interspecies transfer (e.g. in the *Debaryomyces* species), the predominant pattern is one of vertical transmission.

**A.** *Wickerhamiella hasegawae* toxin candidate L0a (plasmid p128, PHY070). β-Estradiol induction. Non-Functional.

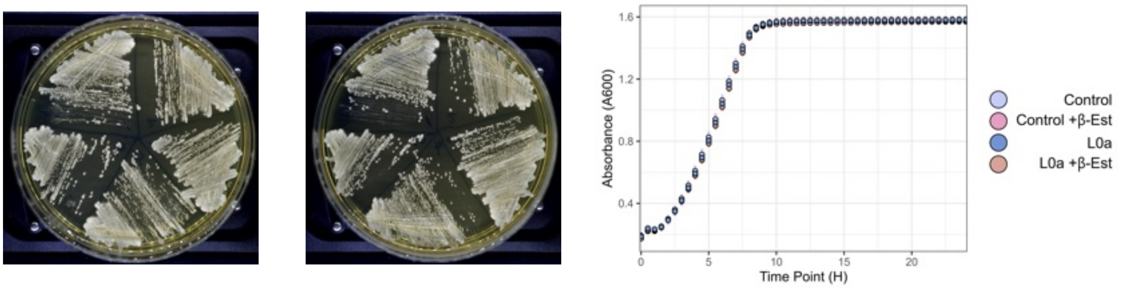


**B.** *Wickerhamiella hasegawae* toxin candidate L0b (plasmid p58, PHY067). β-Estradiol induction. Functional.

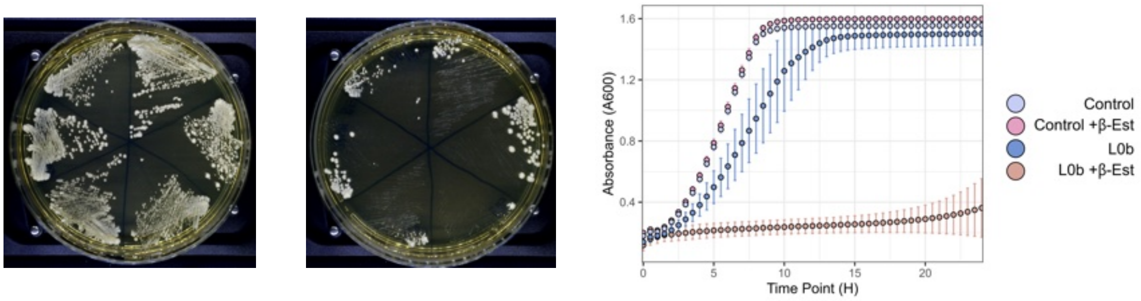


**C.** *Starmerella apicola* toxin candidate L0c (plasmid p61, strain PHY060). β-Estradiol induction. Non-Functional.

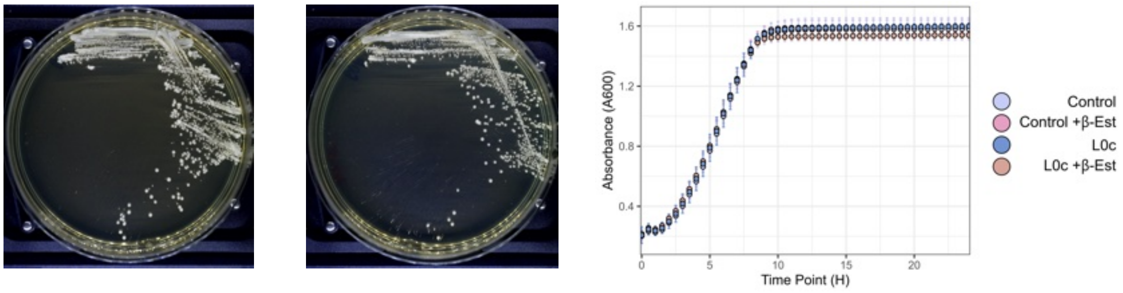


**D.** *Blastobotrys proliferans* toxin candidate L0d (plasmid p92, strain PHY053). β-Estradiol induction. Functional.

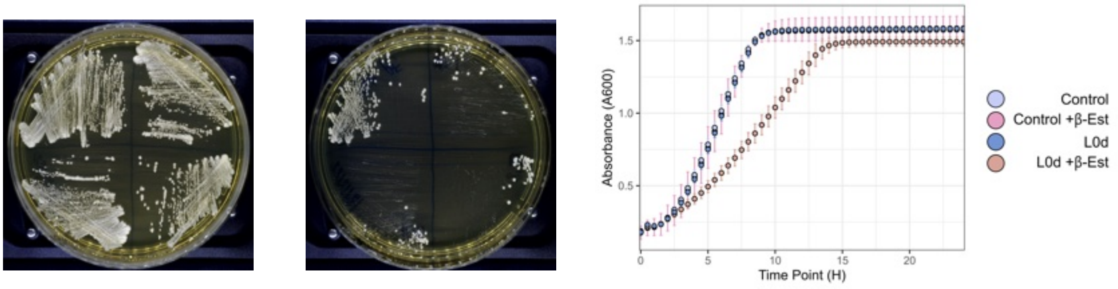


**Figure S2 (page 1).** Functional assays of toxin candidates on solid and liquid media. Each panel shows two plates, YPD uninduced (left) and YPD induced (right) for multiple independent clones of a toxin, and OD600 for growth in liquid media over 24 h with and without toxin induction. Strains were induced with either β-estradiol or galactose, as indicated. See Materials and Methods for details.

**E.** *Kluyveromyces lactis* zymocin L1a (plasmid pPGAL-Z, strain PHY101). Galactose induction. Functional.

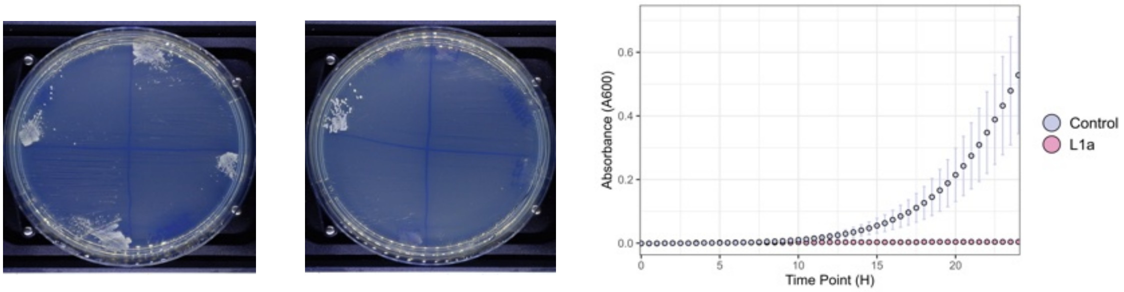


**F-1.** *Lachancea kluyveri* toxin candidate L1b (plasmid p55, strain PHY046). β-Estradiol induction. Functional. Image is also shown in Fig. 4.

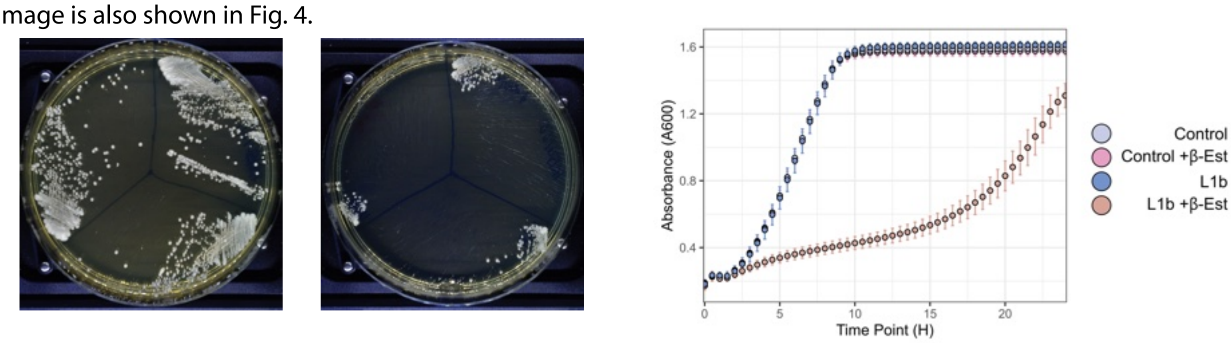


**F-2.** *Lachancea kluyveri* toxin candidate L1b (plasmid pPGAL-55, strain PHY108). Galactose induction. Functional.

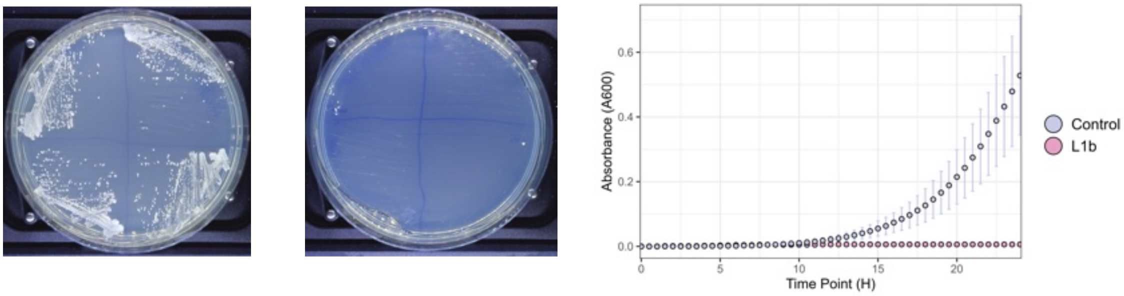


**G.** *Tetrapisispora namnaoensis* toxin candidate L1c (plasmid p129, strain PHY069). β-Estradiol induction. Functional. Image is also shown in Fig. 4.

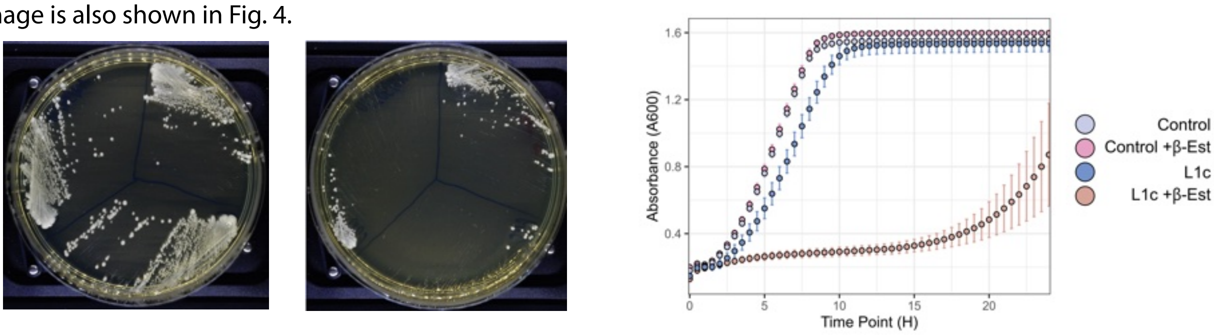


**H.** *Torulaspora delbrueckii* toxin candidate L1d (plasmid p93, strain PHY066). β-Estradiol induction. Functional.

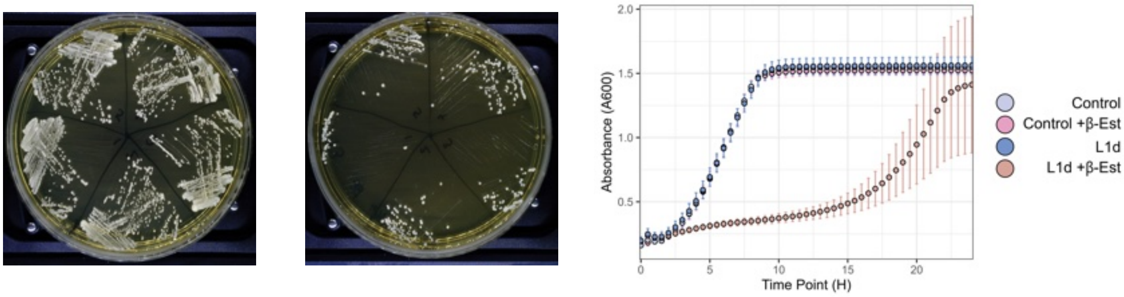


**Figure S2 (page 2).**

**I.** *Torulaspora delbrueckii* toxin candidate L1e (plasmid p89, strain PHY058). β-Estradiol induction. Functional.

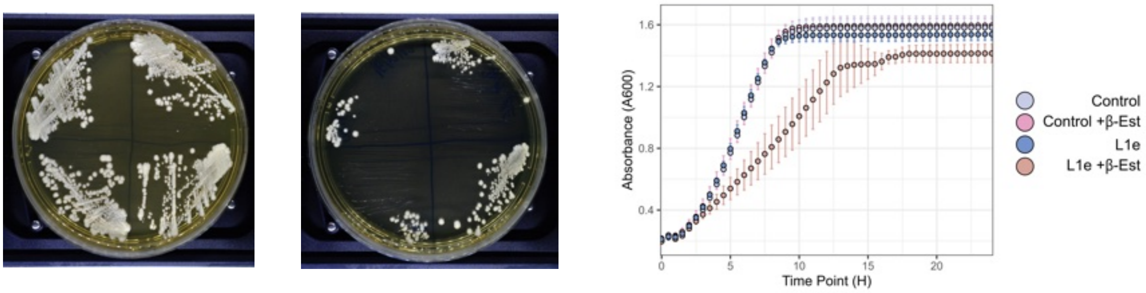


**J.** *Martiniozyma abietophila* toxin candidate L2a (plasmid p67, strain PHY062). β-Estradiol induction. Non-Functional.

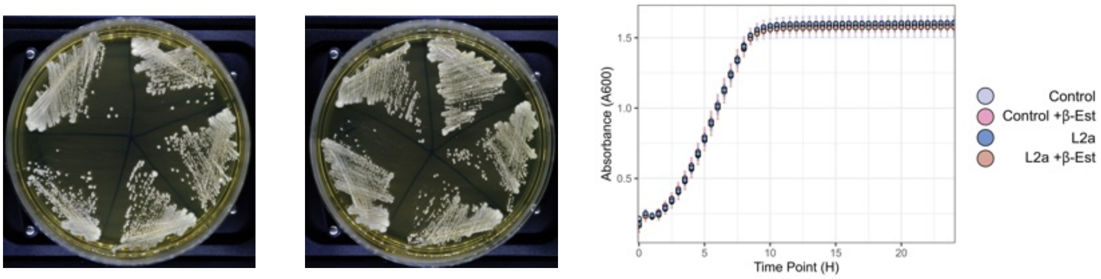


**K.** *Candida boidinii* toxin candidate L2b (plasmid p69, strain PHY050). β-Estradiol induction. Non-Functional.

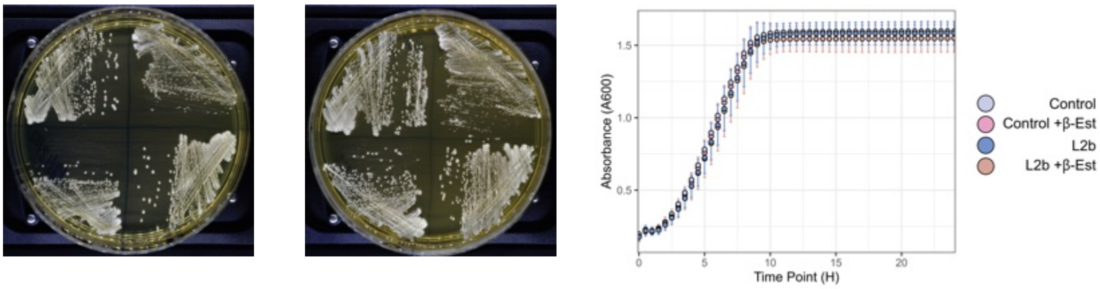


**L.** *Candida inconspicua* toxin candidate L2c (plasmid p76, strain PHY052). β-Estradiol induction. Non-functional.

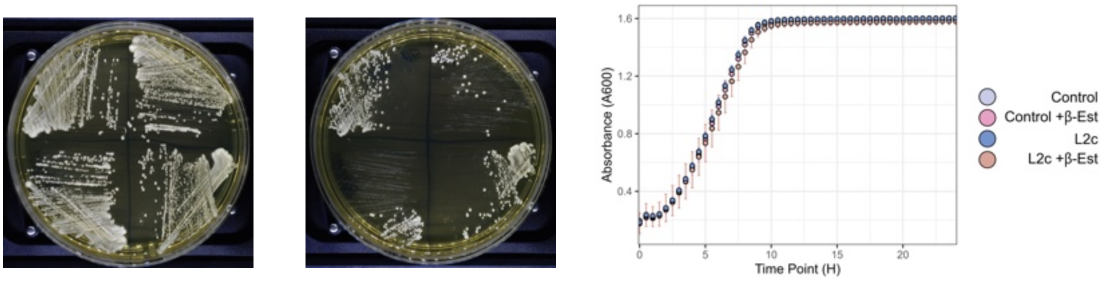


**Figure S2 (page 3).**

**M-1.** *Babjeviella inositovora* S1a (corrected sequence of PiT) (plasmid pPGAL1-140, strain PHY109). Galactose induction. Functional.

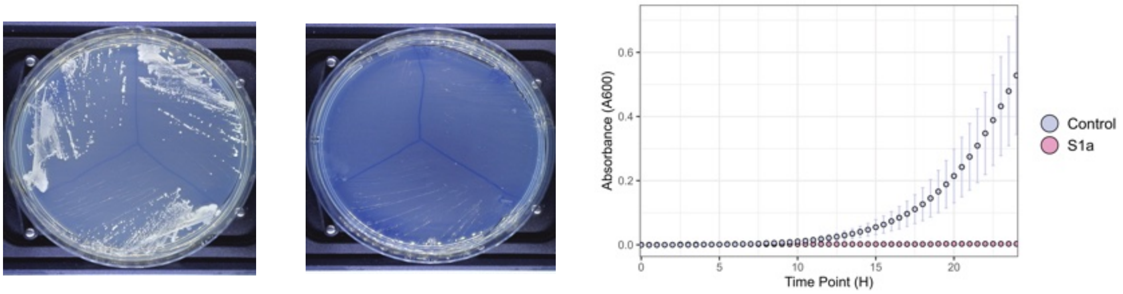


**M-2.** *Babjeviella inositovora* PiT original reported sequence incl. frameshift. (plasmid p71, strain PHY054). β-estradiol induction. Non-Functional.

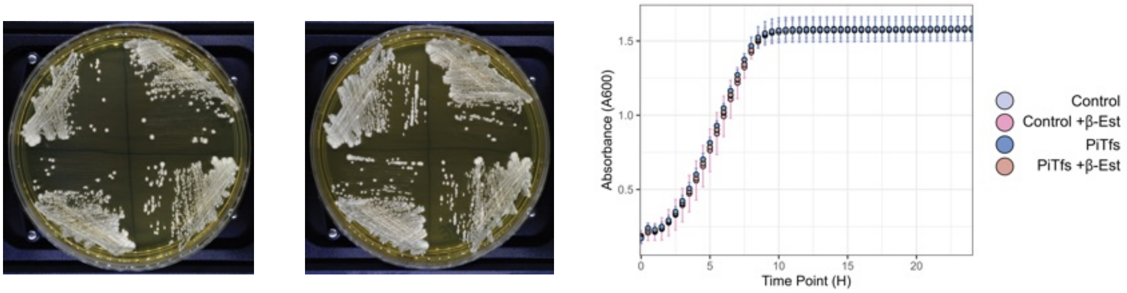


**N-1.** *Millerozyma acaciae* PaT S1c (plasmid p82, strain PHY042). β-estradiol induction. Functional. Image is also shown in Fig. 4.

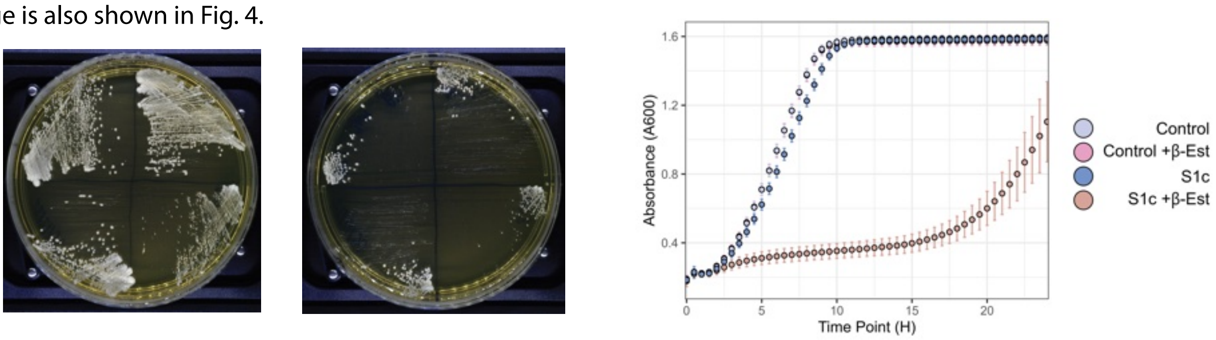


**N-2.** *Millerozyma acaciae* PaT S1c (plasmid pPGAL-82, strain PHY102). Galactose induction. Functional.

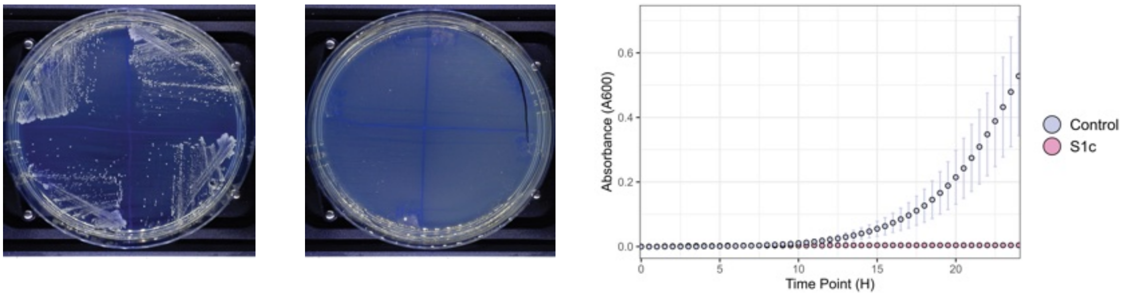


**Figure S2 (page 4).**

**O.** *Metschnikowia lacustris* UWOPS 12-619.2 toxin candidate S1d (plasmid p84, strain PHY064). β-estradiol induction. Functional.

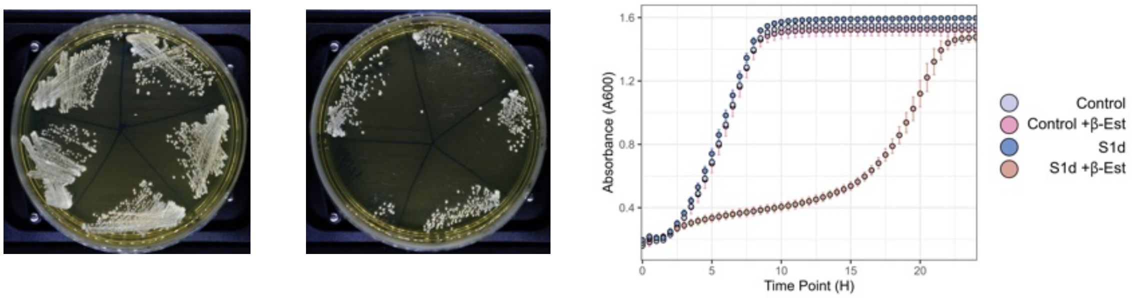


**P.** *Metschnikowia hibisci* toxin candidate S1e (plasmid p62, strain PHY061). β-estradiol induction. Non-Functional.

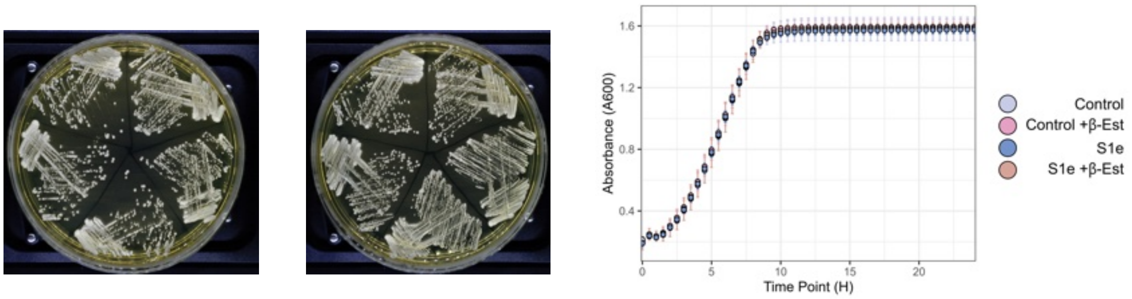


**Q.** *Hyphopichia rhagii* toxin candidate S1g (plasmid p68, strain PHY049). β-estradiol induction. Non-Functional.

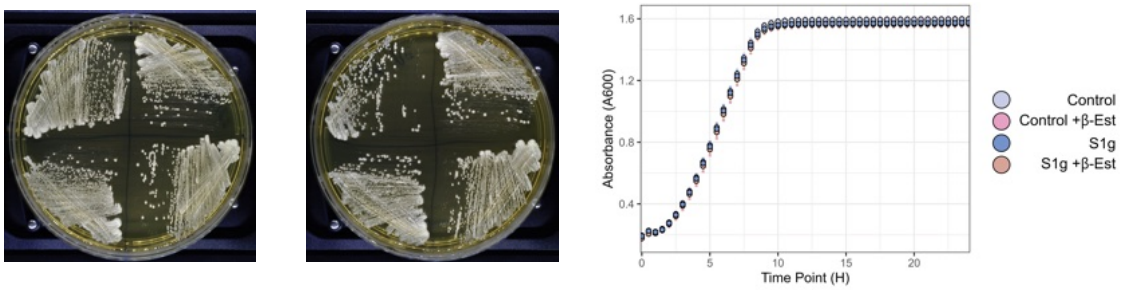


**R.** *Hypopichia rhagii* toxin candidate S1h (plasmid p77, strain PHY063). β-estradiol induction. Functional.

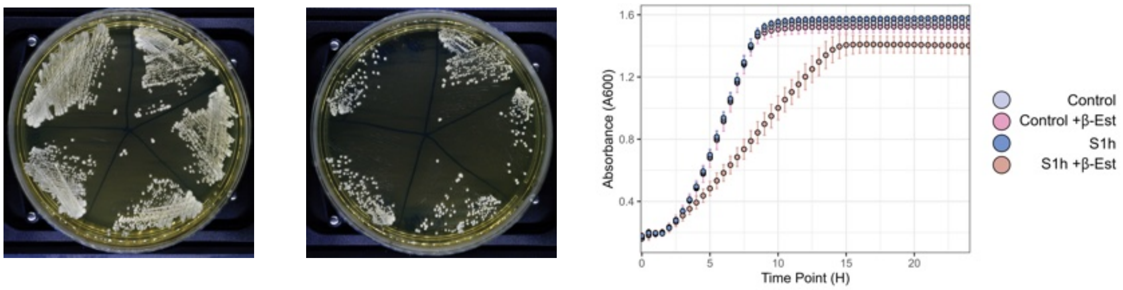


**S.** *Candida gorgasii* toxin candidate S1i (plasmid p70, strain PHY051). β-estradiol induction. Functional.

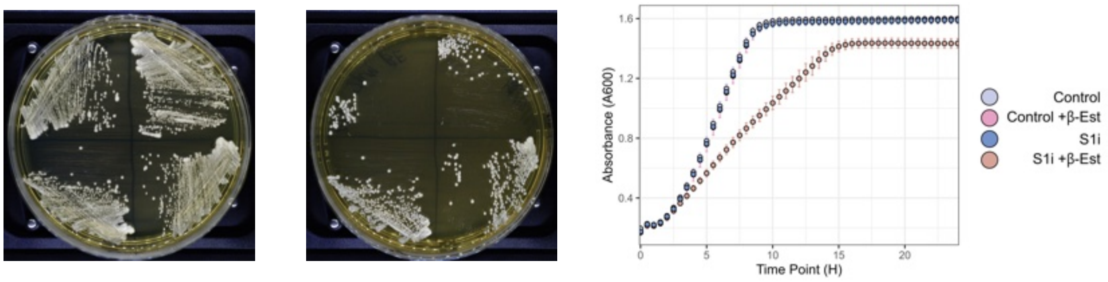


**Figure S2 (page 5).**

**T.** *Spathaspora* sp. JA1 toxin candidate S1j (plasmid p85, strain PHY048). β-estradiol induction. Functional.

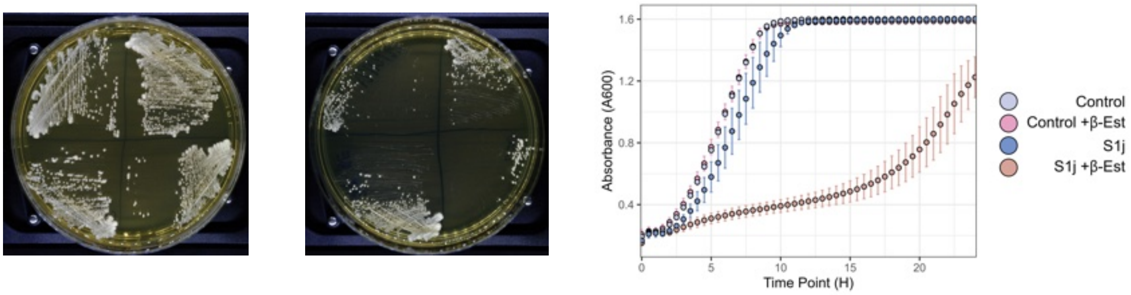


**U.** *Spathaspora girioi* toxin candidate S1k (plasmid p86, strain PHY056). β-estradiol induction. Functional.

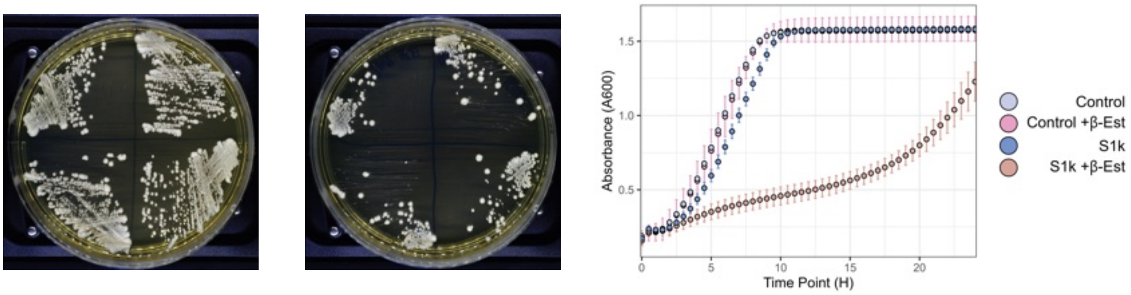


**V.** *Spathaspora girioi* toxin candidate S1l (plasmid p87, strain PHY065). β-estradiol induction. Functional.

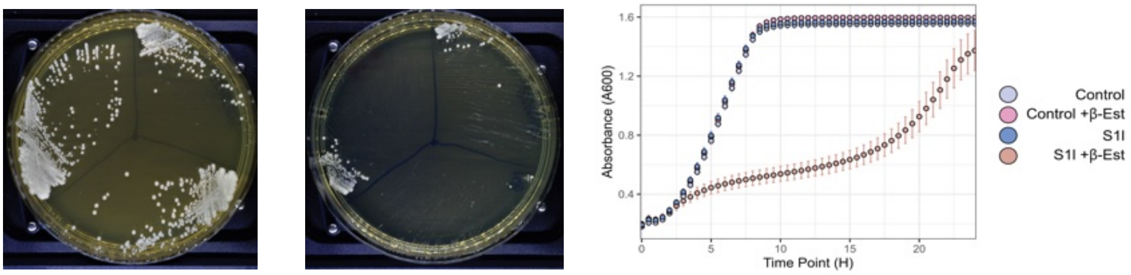


**W.** *Spathaspora girioi* toxin candidate S1m (plasmid p88, strain PHY057). β-estradiol induction. Functional.

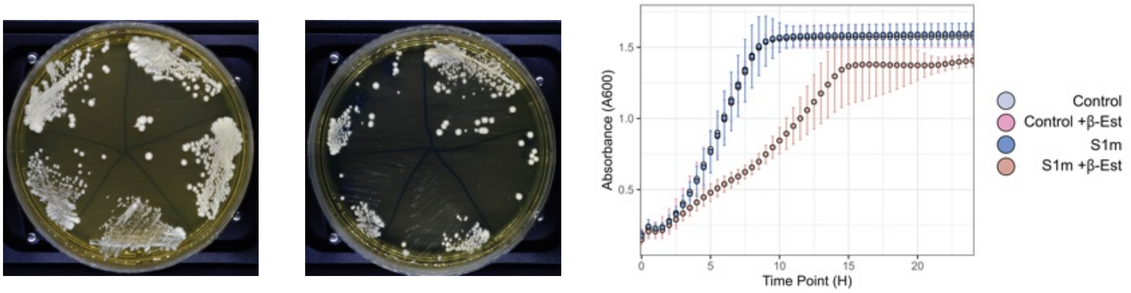


**Figure S2 (page 6).**

**X.** *Debaryomyces robertsiae* toxin candidate S1n (DrT-like) (plasmid pPGAL1-94, strain PHY105). Galactose induction. Functional.

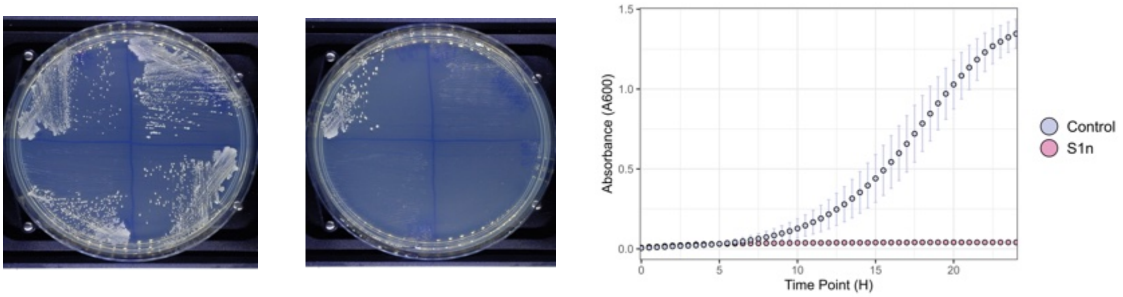


**Y-1.** *Debaryomyces robertsiae* S1o (corrected sequence of DrT)
(plasmid pPGAL1-90, strain PHY103). Galactose induction. Functional.

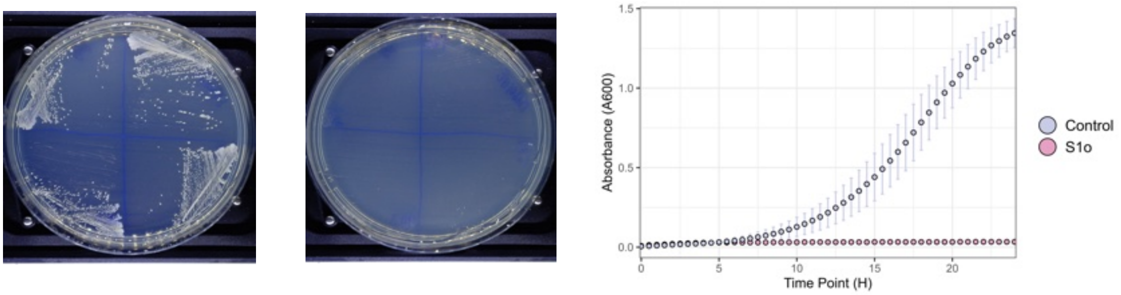


**Y-2.** *Debaryomyces robertsiae* DrT original reported sequence incl. frameshift.
(plasmid pPGAL1-91, strain PHY104). Galactose induction. Non-Functional.

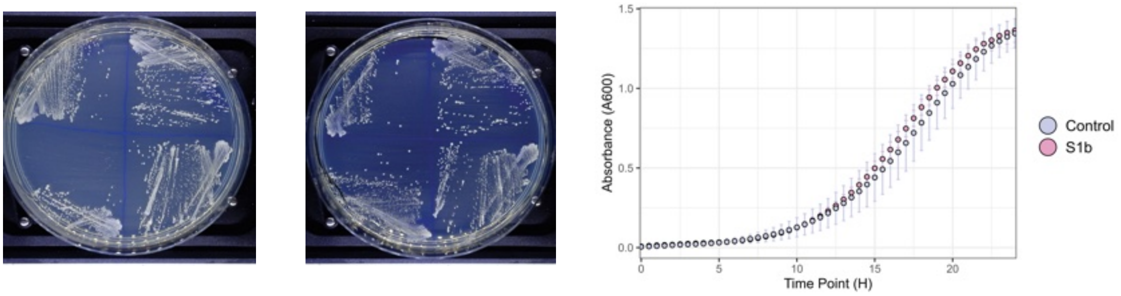


**Y-3.** *Debaryomyces robertsiae* DrT original reported sequence incl. frameshift.
(plasmid p91, strain PHY059). β-estradiol induction. Non-Functional.

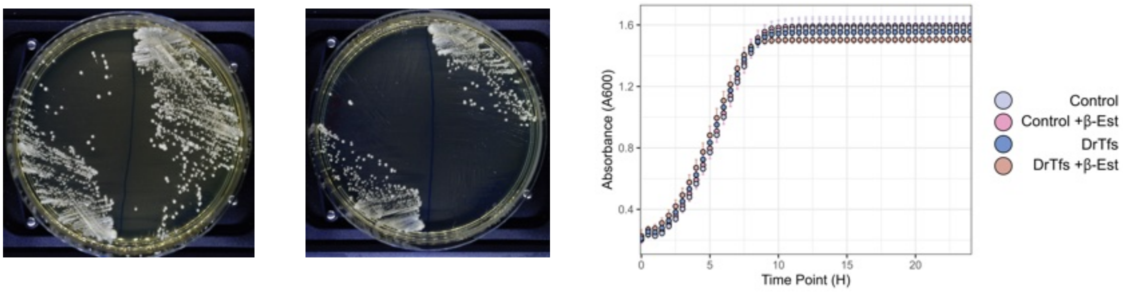


**Figure S2 (page 7).**

**Z.** *Candida oceani* toxin candidate S1q (plasmid p130, PHY068). β-estradiol induction. Non-Functional.

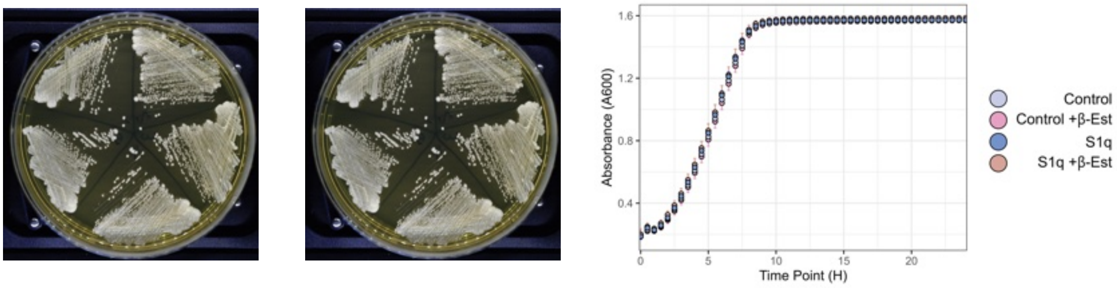


**ZA.** *Candida oceani* toxin candidate S1r (plasmid p131, PHY073). β-estradiol induction. Functional.

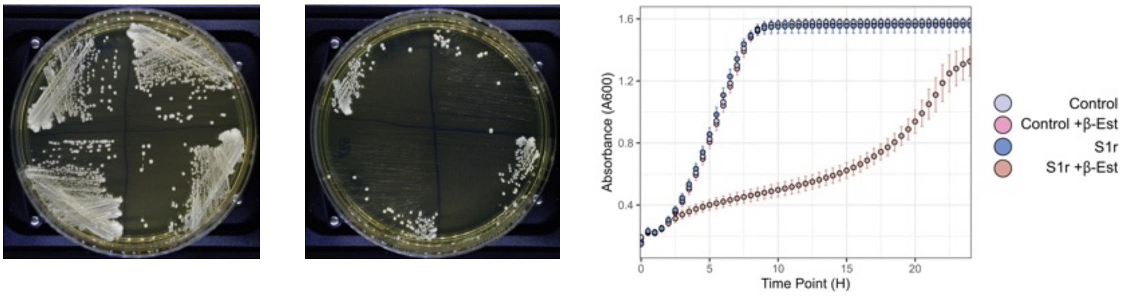


**ZB.** *Saccharomycopsis crategensis* toxin candidate S2a including frameshift correction. (plasmid p75, strain PHY043). β-Estradiol induction. Non-Functional.

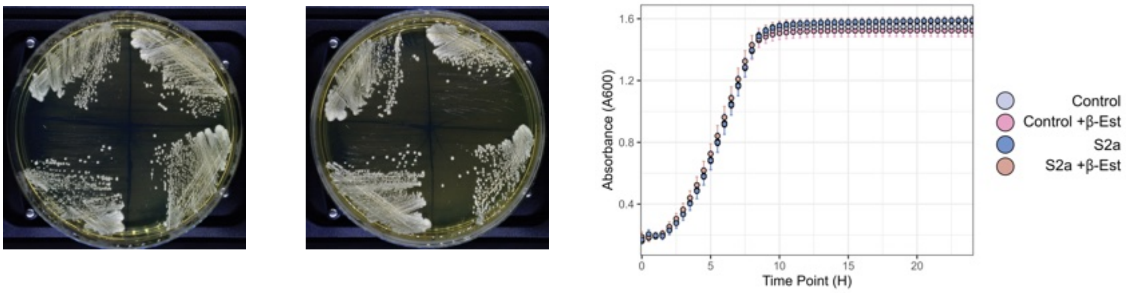


**ZC.** *Saccharomycopsis praedatoria* toxin candidate S2e (plasmid pPGAL-150, strain PHY127). Galactose induction. Functional.

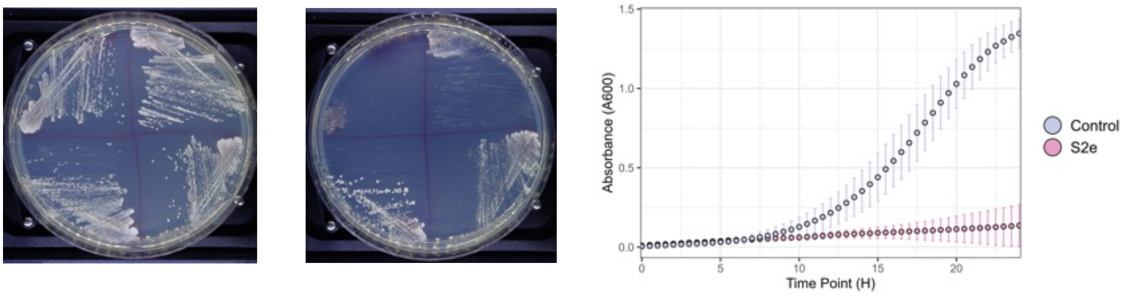


**Figure S2 (page 8).**

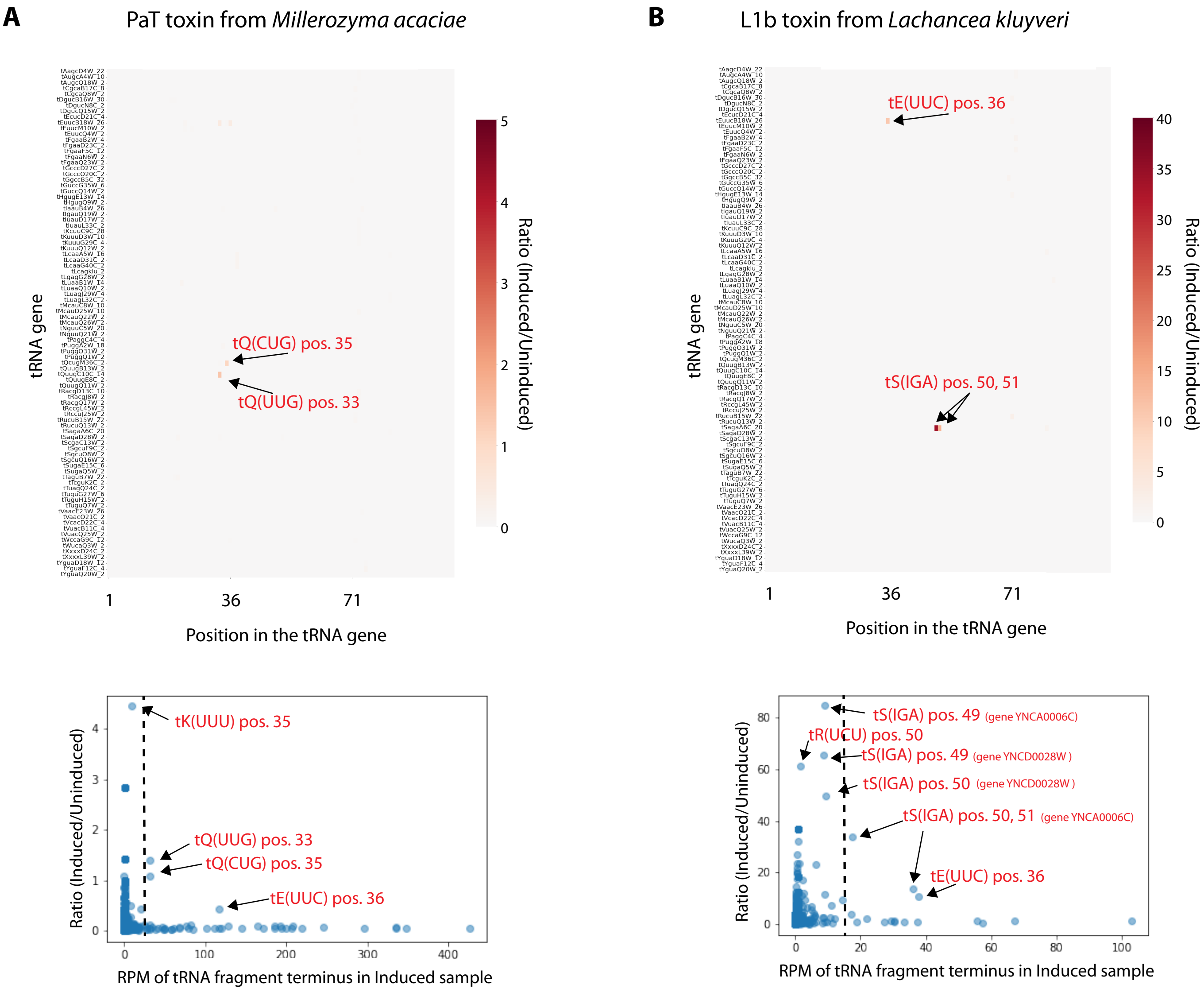


**Figure S3.** cyPhyRNA-seq investigation of tRNA targets of **(A)** the canonical toxin PaT from *Millerozyma acaciae*, and **(B)** the newly discovered toxin L1b from *Lachancea kluyveri*. Sequencing libraries were constructed using *Arabidopsis thaliana* tRNA ligase variant K152A/D726A to capture tRNA fragments having 2′,3′-cP termini, from *S. cerevisiae* strains in which expression of the toxin was either induced or uninduced. The Illumina sequencing reads were aligned to a database consisting of the DNA sequences of every tRNA isodecoder gene in *S. cerevisiae*, to map the tRNA fragment termini. The heatmaps (upper panels) show the locations of fragment termini that are enriched in the induced samples relative to the uninduced samples. Some termini are highlighted with arrows and names indicating the tRNA type and the nucleotide position of the detected terminus within it. The scatterplots (lower panels) illustrate the data processing steps that were applied to remove noise resulting from ratio calculations from low-abundance reads. They compare the ratio of reads in the induced to uninduced samples (Y-axis), to the abundance of the reads in the induced samples (X-axis; RPM, reads per million), for all tRNA fragment termini detected. We applied filters to remove tRNA fragment termini with <1 RPM, and show the ratios in the most abundant 15% of the remaining termini (points to the right of the dashed lines) in the heatmaps.

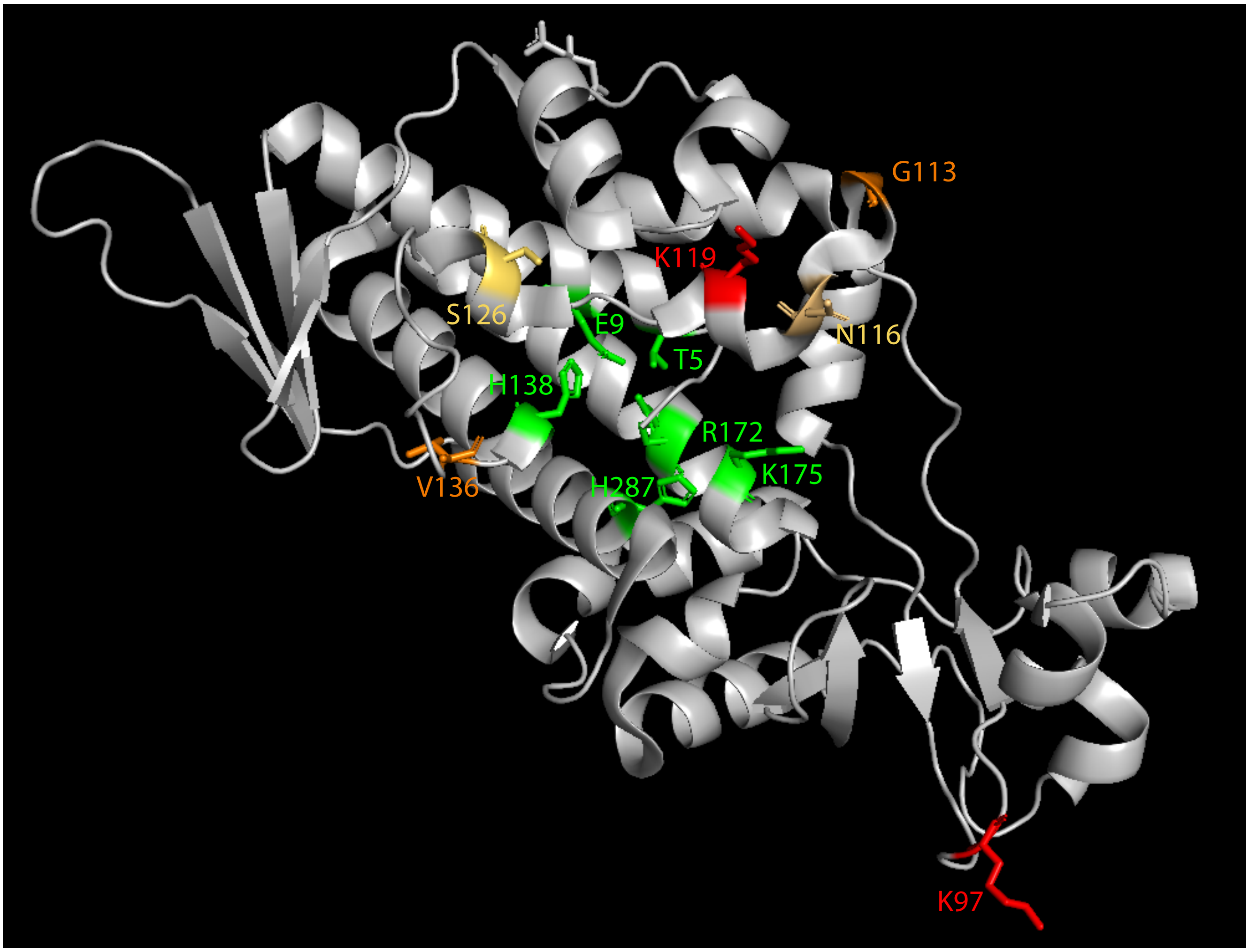


**Figure S4.** Residues of interest highlighted on the structure of PaT (ref. (16); PDB accession 4O87). Green, residues shown to be essential for toxin activity (16). Red, positively selected residues identified by both Naive Empirical Bayes (NEB) and Bayes Empirical Bayes (BEB) with P > 0.99. Orange, positively selected residues identified by both NEB and BEB with P > 0.95. Yellow, residues identified only by BEB as positively selected with P > 0.95. The positively selected sites N116, K119, S126, and V136 are located on the periphery of the active site, suggesting a role in stabilizing an interaction between a target and the active site.

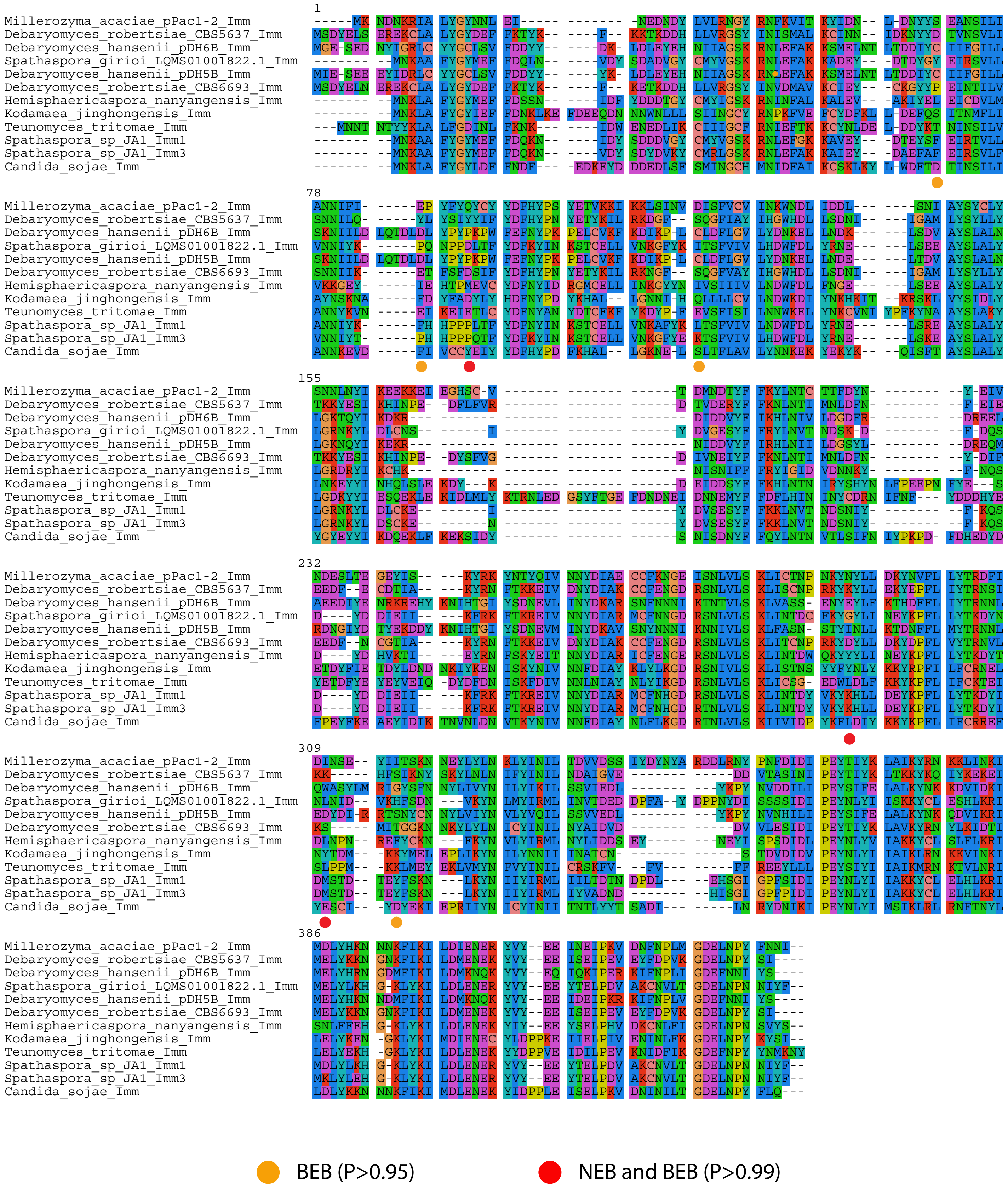


**Figure S5.** Multiple sequence alignment of Imm proteins from the Imm-B cluster, with the positively selected sites identified by CODEML (Table S3) indicated. The three sites identified by both NEB and BEB with P > 0.99 (red dots) correspond to Q68, N220 and I239 of ImmPaT.
